# Resistance mechanisms of SARS-CoV-2 3CLpro to the non-covalent inhibitor WU-04

**DOI:** 10.1101/2023.11.01.564972

**Authors:** Lijing Zhang, Xuping Xie, Hannan Luo, Hongtao Yu, Jing Huang, Pei-Yong Shi, Qi Hu

**Affiliations:** Zhejiang University, Hangzhou, Zhejiang, 310058, China; Westlake AI Therapeutics Lab, Westlake Laboratory of Life Sciences and Biomedicine; Key Laboratory of Structural Biology of Zhejiang Province, School of Life Sciences, Westlake University; Institute of Biology, Westlake Institute for Advanced Study, Hangzhou, Zhejiang, 310024, China; Department of Biochemistry and Molecular Biology, University of Texas Medical Branch, Galveston, TX, 77555, USA

## Abstract

Drug resistance poses a significant challenge in the development of effective therapies against SARS-CoV-2. Here, we identified two double mutations, M49K/M165V and M49K/S301P, in the 3C-like protease (3CLpro) that confer resistance to a novel non-covalent inhibitor, WU-04. Crystallographic analysis indicates that the M49K mutation destabilizes the WU-04 binding pocket, impacting the binding of WU-04 more significantly than the binding of 3CLpro substrates. The M165V mutation directly interferes with WU-04 binding. The S301P mutation, which is far from the WU-04 binding pocket, indirectly affects WU-04 binding by restricting the rotation of 3CLpro’s C-terminal tail and impeding 3CLpro dimerization. We further explored 3CLpro mutations that confer resistance to two clinically used inhibitors: ensitrelvir and nirmatrelvir, and revealed a trade-off between the catalytic activity, thermostability, and drug resistance of 3CLpro. We found that mutations at the same residue (M49) can have distinct effects on the 3CLpro inhibitors, highlighting the importance of developing multiple antiviral agents with different skeletons for fighting SARS-CoV-2. These findings enhance our understanding of SARS-CoV-2 resistance mechanisms and inform the development of effective therapeutics.

## INTRODUCTION

The coronavirus 3C-like protease (3CLpro), also known as main protease (Mpro), plays a crucial role in processing two polyproteins (pp1a and pp1ab) encoded by the virus RNA genome (*1*). Inhibiting the catalytic activity of 3CLpro has been proven to be an effective strategy to block coronavirus replication. Since the emergence of coronavirus disease 2019 (COVID-19) (*2, 3*), caused by the SARS-CoV-2 virus, substantial efforts have been dedicated to the development of SARS-CoV-2 3CLpro inhibitors. Several 3CLpro inhibitors have been approved for treating COVID-19 patients, including the covalent inhibitor PF-07321332 (nirmatrelvir) (*4*) and its analogs SIM0417 (simnotrelvir) and RAY1216 (leritrelvir), as well as the non-covalent inhibitor S-217622 (ensitrelvir) (*5*).

With the increasing clinical use of 3CLpro inhibitors, the emergence of drug resistance has become a growing concern. Although no SARS-CoV-2 variants resistant to 3CLpro inhibitors have been reported in patients to date, several mutations in 3CLpro conferring resistance to nirmatrelvir have been identified through *in vitro* studies (*6–17*). According to the crystal structure of the 3CLpro/nirmatrelvir complex (PDB ID code 7RFS) (*4*), most of these mutation sites are located at three segments within 5 Å of nirmatrelvir, including residues 140–144, 163– 168, and 186–192. Mutations in these segments either directly disrupt their interactions with nirmatrelvir or alter the conformation of the nirmatrelvir-binding pocket, thus leading to drug resistance (*16, 18*). There are also mutations that target residues located far away from nirmatrelvir, such as T21I, P252L, and T304I. Although each of these mutations contributes a modest level of resistance, they are thought to act as initial mutations that facilitate the emergence of additional ones, leading to robust resistance to nirmatrelvir. However, the precise mechanisms by which these mutations confer resistance remain to be elucidated (*8*).

All clinically approved inhibitors of SARS-CoV-2 3CLpro are designed to target the substrate binding pocket of 3CLpro, which raises concern about cross-resistance. Many of the nirmatrelvir-resistant mutations also confer resistance to ensitrelvir (*14–17*). However, mutations confer resistance to one drug but not the other have also been reported. For example, the A173V mutation significantly reduces the potency of nirmatrelvir, but has minimal impact on the potency of ensitrelvir (*15*). In contrast, mutations at M49, such as M49I and M49L, show little impact on the potency of nirmatrelvir but result in substantial resistance to ensitrelvir (*15, 16*). Understanding the spectrum of drug resistance presented by 3CLpro inhibitors that have varying scaffolds and binding modes is crucial for addressing potential cross-resistance issues.

We recently reported the development of a novel class of non-covalent inhibitors targeting coronavirus 3CLpro (*19*). Among them, WU-04 demonstrated significant potency towards 3CLpro in the original SARS-CoV-2 strain (wild-type, WT) and the Omicron variants. It also effectively inhibited other coronaviruses, such as SARS-CoV and MERS-CoV (*19*). Now, WU- 04 is undergoing clinical trials for the treatment of COVID-19. In this study, we have identified mutations in SARS-CoV-2 3CLpro that confer resistance to WU-04. By determining their crystal structures, we have elucidated the mechanisms for WU-04 resistance. Additionally, we studied mutations that confer resistance to ensitrelvir and nirmatrelvir, and assessed the cross-resistance of 3CLpro carrying these mutations to WU-04, ensitrelvir and nirmatrelvir.

## RESULTS

### Mutations in 3CLpro Confer SARS-CoV-2 resistance to WU-04

Having identified the potent anti-SARS-CoV-2 inhibitor WU-04, we sought to explore the possibility of developing WU-04-resistant mutations in SARS-CoV-2. Selection of the resistant virus was performed in African green monkey kidney epithelial Vero E6 cells using a reporter SARS-CoV-2 (this virus showed significantly reduced virulence *in vivo* compared to the wild-type SARS-CoV-2). WU-04 blocked SARS-CoV-2 replication in Vero E6 cells with an IC_50_ of 10–20 nM. By serially passaging the reporter SARS-CoV-2 in Vero E6 cells in the presence of increasing concentrations of WU-04, we identified four strains of SARS-CoV-2 that could survive 10 µM WU-04 (Fig. S1A). Two strains carried double nucleotide mutations T10200A (3CLpro: M49K) and A10547G (3CLpro: M165V) in the viral genomes, whereas the other two strains carried double mutations T10200A (3CLpro: M49K) and T10955C (3CLpro: S301P). Fortunately, the three drug-resistance mutations were found in the GISAID database (https://gisaid.org/) with a relatively low frequency (Table S1).

In the crystal structure of the SARS-CoV-2 3CLpro/WU-04 complex, residues M49 and M165 pack against the bromophenyl ring of WU-04, suggesting that these mutations directly hinder the binding of WU-04 to 3CLpro (Fig. 1A). The identification of resistant mutations in the WU-04 binding pocket of 3CLpro validates the on-target effect of WU-04 in cellular assays. Residue S301 is located at the end of the last α-helix of 3CLpro. It precedes the C-terminal tail (residues 301–306), which is involved in 3CLpro homodimerization (Fig. 1A). The S301P mutation may indirectly impede WU-4 binding through modulating the C-terminal tail of 3CLpro.

**Fig. 1.**
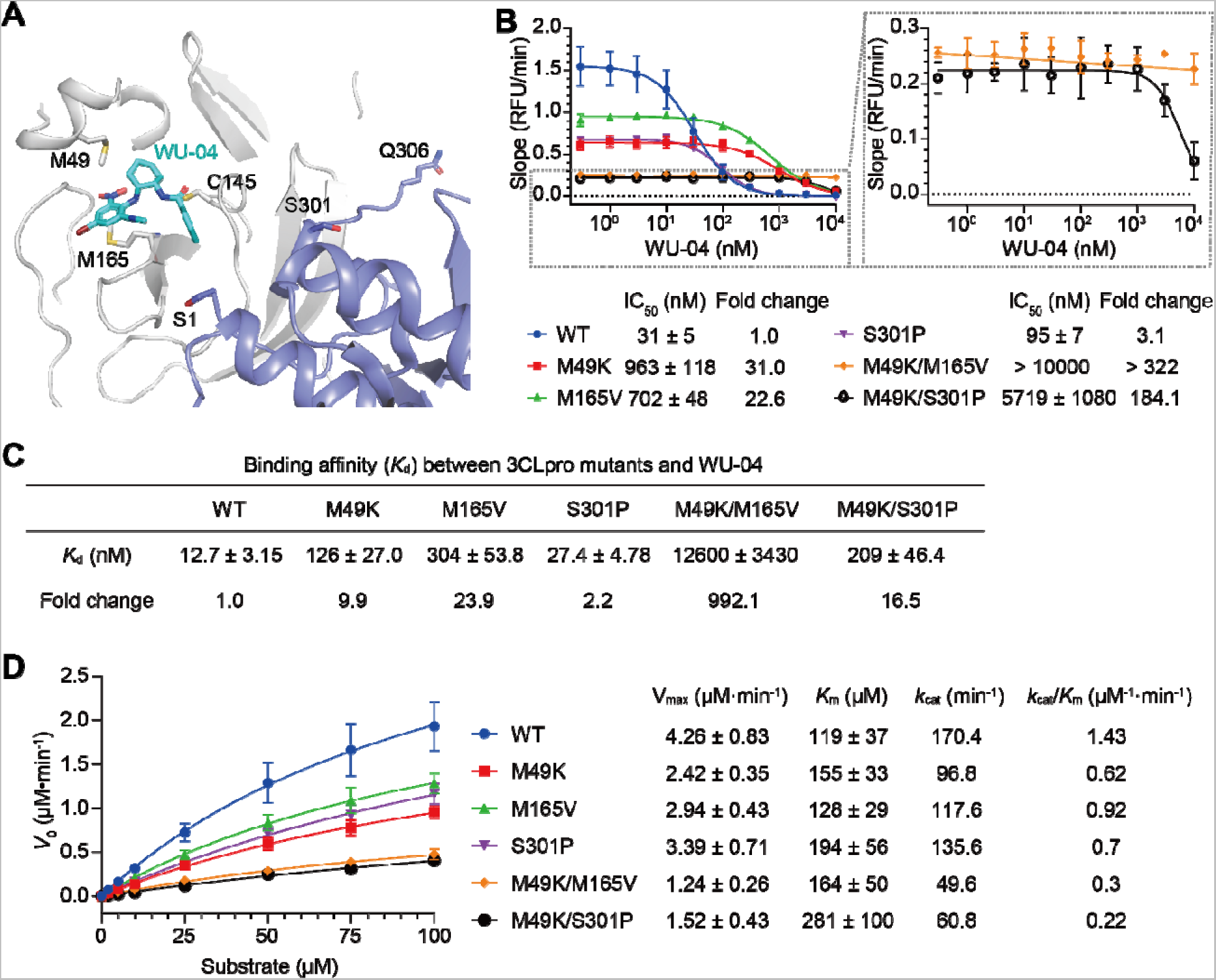
Characterization of WU-04-resistant mutations in SARS-CoV-2. (**A**) Two double mutations M49K/M165V and M49K/S301P of SARS-CoV-2 3CLpro were identified as WU-04-resistant mutations. Residues M49, M165, and the catalytic residue C145 from one 3CLpro protomer, along with residue S301 and the N- and C-terminal residues S1 and Q306 from the other 3CLpro protomer within the same 3CLpro homodimer, are depicted as sticks in the crystal structure of the 3CLpro/WU-04 complex (PDB ID code 7EN8). (**B**) The inhibitory activity of WU-04 against the SARS-CoV-2 3CLpro mutants was assessed using a fluorescence resonance energy transfer (FRET)-based assay. The two double mutants M49K/M165V and M49K/S301P showed the strongest resistance to WU-04. The data represents the mean ± SD of three independent measurements. (**C**) The binding affinity (*K*_d_) between WU-04 and each 3CLpro mutant was measured using isothermal titration calorimetry (ITC), and then normalized to that between WU-04 and the wild-type 3CLpro to get the number of fold change. (**D**) The catalytic activity of each 3CLpro mutant was evaluated using the FRET-based assay. The data represent the mean ± SD of three independent measurements

We purified the WT 3CLpro and the WU-04-resistant mutants and measured their inhibition by WU-04 using a fluorescence-based enzyme assay (Fig. 1B). Both the M49K and M165V mutations increased the IC_50_ of WU-04 against 3CLpro by more than 20-fold. Combination of the two mutations further increased the IC_50_ to greater than 10 µM. The S301P mutation alone slightly increased the IC_50_, but in combination with the M49K mutation, drastically increased the IC_50_ to greater than 5 µM.

We then measured the binding affinities of these 3CLpro mutants to WU-04 using isothermal titration calorimetry (ITC). Consistent with the decreased sensitivity of these mutants to WU-04, the dissociation constants (*K*_d_) for these mutants bound to WU-04 were all increased (Fig. 1C and Fig. S1B). Interestingly, though the M49K/S301P mutant showed stronger WU-04 resistance than the M165V mutant, its binding affinity to WU-04 was higher than that of the M165V mutant, indicating a resistant mechanism beyond the decreased WU-04 binding.

We next evaluated the effects of these mutations on the catalytic activity of SARS-CoV-2 3CLpro (Fig. 1D). The protease activities of each mutant at different substrate concentrations were measured to obtain the *V*_max_, *K*_m_ and *k*_cat_, and calculate the *k*_cat_/*K*_m_ ratio as a measurement of the catalytic activity. All these mutants showed higher *K*_m_ and smaller *k*_cat_ value in comparison with the WT 3CLpro. Specifically, the *k*_cat_/*K*_m_ ratios of the double mutants, M49K/M165V and M49K/S301P, were only about one fifth and one seventh of that of the WT 3CLpro, respectively. These results demonstrate that there is a trade-off between the catalytic activity and the WU-04 resistance of 3CLpro. The reduction in *k*_cat_ could potentially be attributed to either a disturbance within the catalytic site or a destabilization of the 3CLpro dimer.

### The M49K mutation disturbs the substrate binding pocket of 3CLpro

To understand how these mutations affect the WU-04 sensitivity and the catalytic activity of 3CLpro, we solved the crystal structures of these mutants (Table S2) and aligned them with the crystal structure of the WT 3CLpro. The overall structures of these mutants are very similar to that of the WT 3CLpro (PDB ID code 6M03), with RMSD values below 0.4 Å.

In the crystal structure of the M49K mutant, three regions around the substrate binding pocket show conformational changes in comparison with the structure of the WT 3CLpro (Fig. 2A). One is the short helix (residues 45–51) in which the M49K mutation is located. The electron density of this helix in the crystal structure of the M49K mutant is so weak that its structure cannot be modeled (Fig. S2A and B), indicating that the M49K mutation destabilized this short helix. The other two regions are the loop (residues 167–171) after M165 and the linker (residues 187–196) connecting domains II and domain III of 3CLpro. Both show slight conformational changes upon introducing the M49K mutation. Conformational changes in the three regions are also observed in the crystal structure of the M49K/M165V double mutant (Fig. 2B and Fig. S2C), but not in the crystal structure of the M165V mutant (Fig. 2C and Fig. S2D) or that of the S301P mutant (Fig. 2D). These structures demonstrate that the M49K mutation disturbs the local structure around the substrate binding pocket of 3CLpro, but the M165V mutation does not induce conformational changes.

**Fig. 2.**
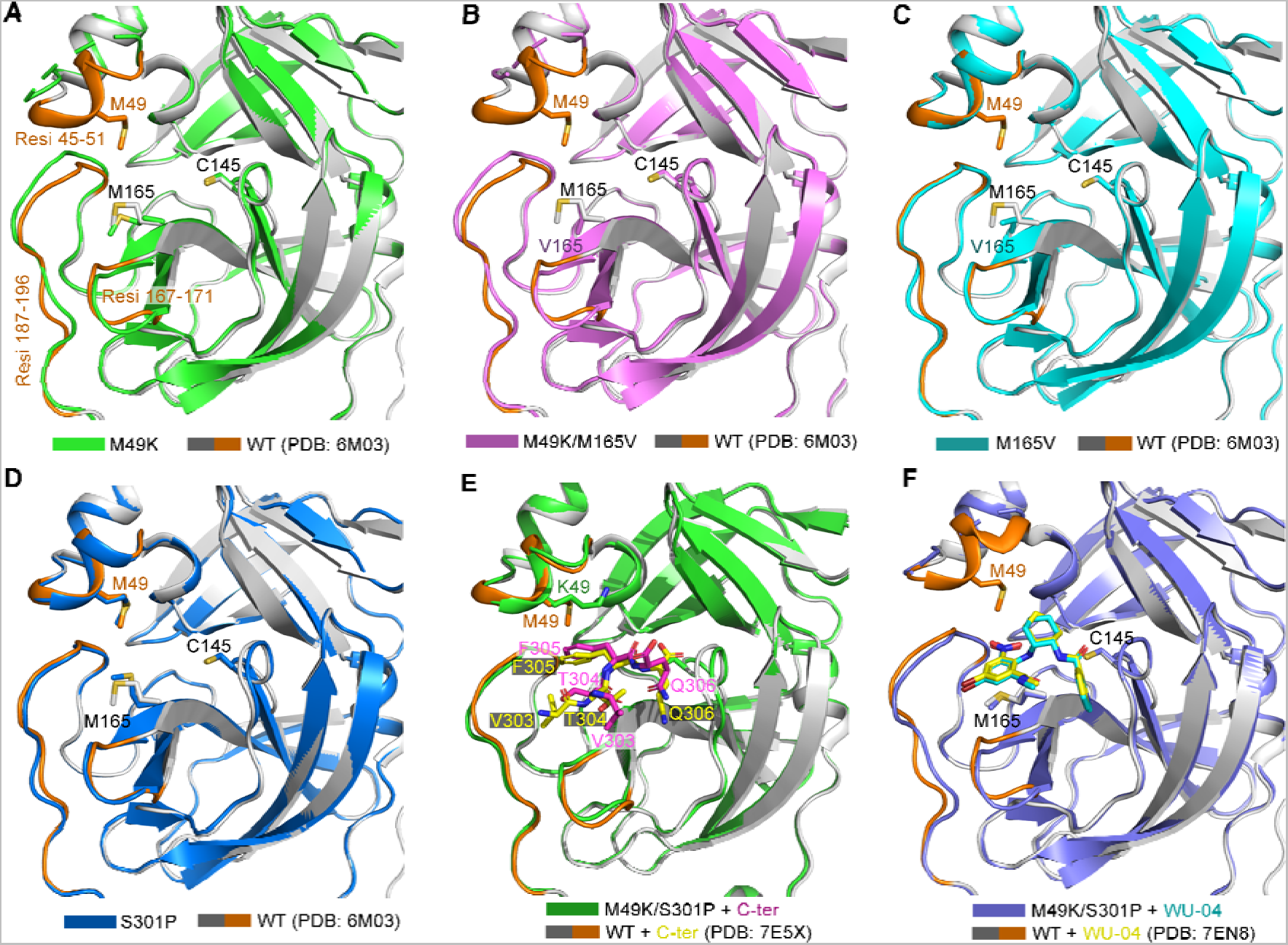
The WU-04 resistant mutation M49K perturbs the substrate binding pocket of SARS-CoV-2 3CLpro. (**A-D**) Alignment of the crystal structures of the WU-04 resistant SARS-CoV-2 3CLpro mutants M49K (**A**), M49K/M165V (**B**), M165V (**C**), and S301P (**D**) with the crystal structure of the WT 3CLpro (PDB ID code 6M03). (**E**) Alignment of the crystal structure of the double mutant M49K/S301P with that of the WT 3CLpro in the post-cleavage state (PDB ID code 7E5X). The C-terminal tail from another molecule of 3CLpro in the M49K/S301P structure and that in the WT 3CLpro structure were shown as sticks and colored magentas and yellow, respectively. (**F**) Alignment of the crystal structure of the double mutant M49K/S301P in complex with WU-04 with the crystal structure of the WT 3CLpro in complex with WU-04 (PDB ID code 7EN8). WU-04 in the M49K/S301P structure and that in the WT 3CLpro structure are colored cyan and yellow, respectively. Three regions in the WT 3CLpro, including a short helix where M49 is located (residues 45–51), a short loop (residues 167–171), and the linker connecting domain II and III of 3CLpro (residues 187–196), are colored brown in the crystal structure.

Unexpectedly, residues 45–51 in the crystal structure of the M49K/S301P double mutant have good electron density (Fig. S2E) and the conformation is similar to that in the structure of the WT 3CLpro. Structural analysis revealed that the C-terminus of one molecule of the M49K/S301P double mutant was docked into the substrate binding pocket of another molecule of this mutant (Fig. 2E), thus this structure may represent the post-cleavage state of the M49K/S301P double mutant. Alignment of this structure with the crystal structure of the WT 3CLpro in the post-cleavage state (PDB ID code 7E5X) shows that the two structures are almost identical to each other, except that residues V303 and T304 from the C-terminus of another molecule of 3CLpro have different orientations in the substrate binding pockets (Fig. 2E). We have also solved the crystal structure of the M49K/S301P double mutant in complex with WU- 04 and found that residues 45–51 have poor electron density (Fig. 2F and Fig. S2F), similar to that in the M49K structure. The conformation and binding position of WU-04 in this structure are the same as that in the crystal structure of the WT 3CLpro/WU-04 complex (PDB ID code 7EN8). These findings suggest that the conformation of residues 45–51 altered by the M49K mutation can be stabilized by binding to 3CLpro substrates but not WU-04. Thus, the M49K mutation has a greater effect on WU-04 binding than on 3CLpro substrate binding.

### The S301P mutation restricts the rotation of the C-terminal tail of 3CLpro

In contrast to the M49K and M165V mutations, which directly affect the catalytic activity of 3CLpro and inhibit 3CLpro binding, the S301P mutation affects a residue located far from the WU-04 binding site. S301 is located at the end of the last helix of 3CLpro. In the crystal structure of the mature WT 3CLpro (PDB ID code 6M03), the C-terminal tail (residues 301–306) of each 3CLpro molecule binds to the other 3CLpro molecule in the same 3CLpro homodimer (Fig. 3A and B), but in the M49K/S301P and S301P structures this C-terminal tail is oriented towards a different direction so that it is no longer involved in 3CLpro dimerization (Fig. 3C, Fig. 3D and Fig. S3A). In the crystal structure of the WT 3CLpro in the post-cleavage state (PDB ID code 7E5X), the C-terminal tails of the two 3CLpro molecules in each 3CLpro homodimer have distinct orientations (Fig. 3E, Fig. S3C and S3D): the one that represents the post-cleavage state of 3CLpro docks into the substrate binding pocket of 3CLpro in another 3CLpro homodimer (chain A), in an orientation similar to that in the M49K/S301P structure; while the other is involved in 3CLpro homo-dimerization (chain B), with its orientation being similar to that in the structure of the mature WT 3CLpro (PDB ID code 6M03). The different orientations are caused by the rotation of the backbone Φ angle of S301 (Fig. 3E). We speculate that, after cleavage, the C-terminal tail of 3CLpro switches from the post-cleavage state (chain A) to the mature state (chain B) to stabilize the 3CLpro homodimer.

**Fig. 3.**
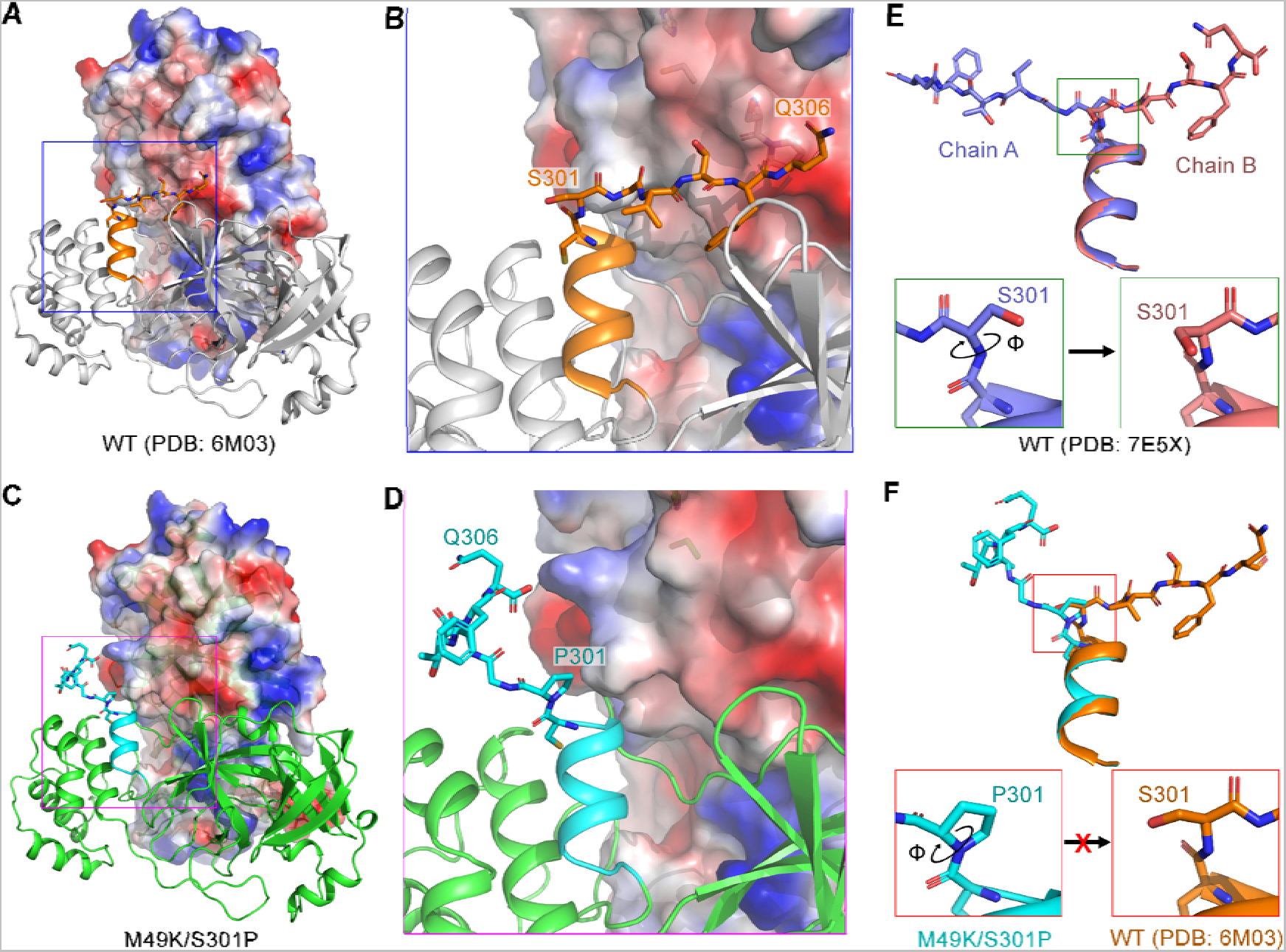
The S301P mutation locks the C-terminal tail of 3CLpro in the post-cleavage state. (**A**, **B**) In the crystal structure of the WT 3CLpro in the mature state (PDB ID code 6M03), the C-terminal tail (residues 301–306, showed in sticks and colored orange) of one 3CLpro protomer binds to the other 3CLpro protomer (the protein contact potential was calculated using PyMOL) within the same 3CLpro homodimer. (**C**, **D**) In the crystal structure of the M49K/S301P double mutant, the C-terminal tail (colored cyan) of one 3CLpro protomer turns away from the other 3CLpro protomer within the same 3CLpro homodimer. (**E**) Alignment of the C-terminal tails of the two 3CLpro protomers (colored purple and salmon, respectively) within the same 3CLpro homodimer in the crystal structure of the WT 3CLpro in the post-cleavage state (PDB ID code 7E5X). The two C-terminal tails have distinct orientations. Rotation of the Φ angle of S301 switches the C-terminal tail from the post-cleavage state (chain A) to the mature state (chain B). (**F**) Alignment of the C-terminal tails of 3CLpro in the M49K/S301P structure (colored cyan) with that in the mature WT 3CLpro structure (colored orange). In the M49K/S301P mutant, the Φ angle of P301 is fixed, therefore, the C-terminal tail of 3CLpro is locked at the post-cleavage state.

When S301 is mutated to a proline residue, the Φ angle of P301 is fixed. Consequently, the C-terminal tail cannot be rotated and is restricted to a conformation that cannot contribute to the homo-dimerization of 3CLpro (Fig. 3F, Fig. S3B, S3E and S3F). This finding indicates that the S301P mutation destabilizes the homodimers of 3CLpro.

### Drug-resistant mutations decrease the catalytic activity and destabilize 3CLpro

We also studied the mutations that make 3CLpro resistant to other two inhibitors: the non-covalent inhibitor ensitrelvir and the covalent inhibitor nirmatrelvir, with the aim of understanding the similarities and differences between these mutations. Initially, we focused on the impact of these mutations on the catalytic activity and thermostability of 3CLpro.

In contrast to WU-04, which occupies the S1, S2 and S4 sites of the substrate binding pocket of 3CLpro (Fig. 4A and D), ensitrelvir occupies the S1, S1’ and S2 sites (Fig. 4B and E). Nirmatrelvir also occupies S1, S2 and S4 (Fig. 4C and F), but the specific interactions with 3CLpro differ from those observed with WU-04. A few mutations in 3CLpro have been reported to confer resistance to ensitrelvir; among them, mutations at M49 of 3CLpro showed the strongest effects (*15, 20*). We chose M49I as a representative mutation. Additionally, we analyzed the crystal structure of 3CLpro in complex with ensitrelvir (PDB ID code 7VU6) and found that the side chain of T25 was in close proximity to the methylindazol ring of ensitrelvir (Fig. 4E). Therefore, we also selected the T25I and T25V mutations as potential ensitrelvir-resistant mutations. For nirmatrelvir, resistance mutations or deletions at nearly all residues within the nirmatrelvir binding pocket have been reported. We selected the mutations and deletions that have been reported to confer strong resistance to nirmatrelvir.

**Fig. 4.**
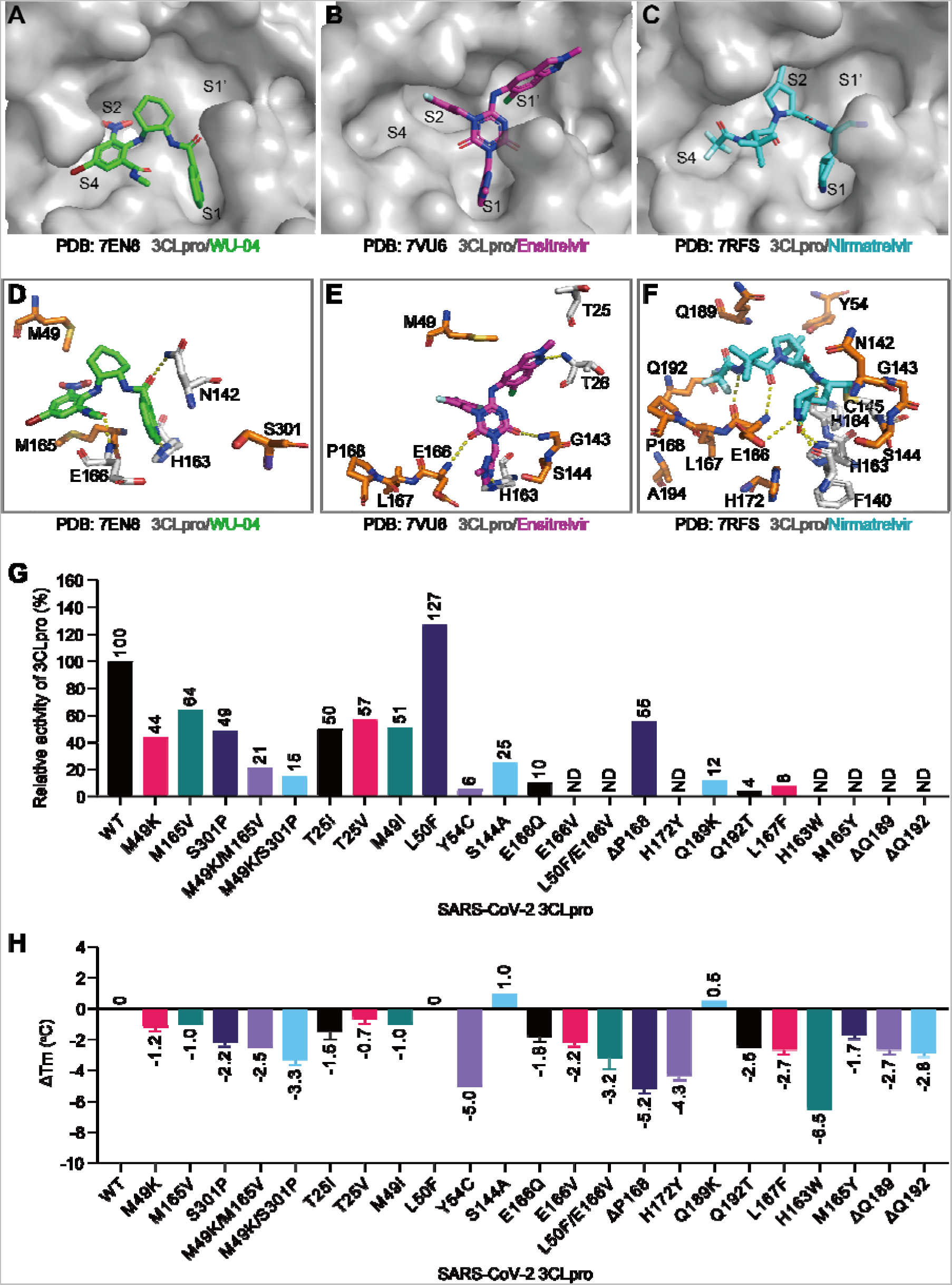
3CLpro mutations that confer resistance to WU-04, ensitrelvir and nirmatrelvir decreased the catalytic activity and thermostability of 3CLpro. (**A-C**) Occupancy of the substrate binding pocket of SARS-CoV-2 3CLpro by the non-covalent inhibitors WU-04 (**A**) and ensitrelvir (**B**), and the covalent inhibitor nirmatrelvir (**C**). (**D-F**) Interactions between SARS-CoV-2 3CLpro and WU-04 (**D**), between SARS-CoV-2 3CLpro and ensitrelvir (**E**), and between SARS-CoV-2 3CLpro and nirmatrelvir (**F**). Hydrogen bonds are indicated by yellow dash lines. The residues mutation of which may confer resistance to the three inhibitors are colored orange. (**G**) Relative catalytic activities of the drug-resistant 3CLpro mutants. The catalytic activity (*k*_cat_/*K*_m_) of each mutant was evaluated using a FRET-based assay with three independent measurements, and then normalized to that of the WT 3CLpro. (**H**) Changes in the melting temperature (T_m_) of SARS-CoV-2 3CLpro induced by drug-resistant mutations. The T_m_ of each mutant was evaluated using a thermal shift assay. The data represent the mean ± SD of technical triplicate.

All mutants were recombinantly expressed in *E. coli* and purified to homogeneity. The *k*_cat_/*K*_m_ ratio of each mutant was calculated and normalized to that of the WT 3CLpro (Fig. 4G and Fig. S4). All these mutants, except L50F, showed decreased catalytic activities. In comparison with 3CLpro mutants that are resistant to the non-covalent inhibitors WU-04 and ensitrelvir, most of the nirmatrelvir-resistant mutants exhibited much lower catalytic activities. For the E166V, L50F/E166V, H172Y, H163W and M165Y mutants, as well as 3CLpro with Q189 or Q192 deletions (ΔQ189 and ΔQ192), their activities in our assay were so low that their *V*_max_ and *K*_m_ values could not be determined and were therefore labeled as not detectable (ND).

It is notable that the L50F mutant demonstrated higher activity as compared to the WT 3CLpro: the *k*_cat_/*K*_m_ ratio of the L50F mutant was 1.82 µM^-1^·min^-1^, higher than that of the WT 3CLpro (1.43 µM^-1^·min^-1^). This observation is distinct from that in previous studies in which the catalytic activity of the L50F mutant was reported as only 0.1% or 0.4% of that of the WT 3CLpro (*7, 9*). The L50F mutation also enhanced the catalytic activity of the El66V mutant (Fig. S4C).

We also evaluated the impact of these drug-resistant mutations on the thermostability of 3CLpro by using a thermal shift assay (*21*). Nearly all these mutations or deletions resulted in a decrease in the melting temperature (T_m_) of 3CLpro (Fig. 4H and Fig. S5). Among them, the Y54C mutation, P168 deletion (ΔP168), and H163W mutation decreased the T_m_ of 3CLpro by a minimum of 5 °C. In contrast, three mutations — L50F, S144A, and Q189K — displayed no effect or led to a slight increase in the T_m_ of 3CLpro.

### Cross-resistance of purified 3CLpro mutants to non-covalent and covalent inhibitors

We subsequently assessed the inhibitory activities of WU-04, ensitrelvir, and nirmatrelvir against purified 3CLpro using the fluorescence-based enzyme assay. For the non-covalent inhibitors WU-04 and ensitrelvir, their IC_50_ values against each mutant were determined (Fig. S6) and normalized to the IC_50_ values against the WT 3CLpro (Fig. 5A). For the covalent inhibitor nirmatrelvir, the inhibition constant (*K*_i_) against each mutant was calculated (Fig. S7), following the method described previously (*4*), and normalized to that against the WT 3CLpro (Fig. 5A).

**Fig. 5.**
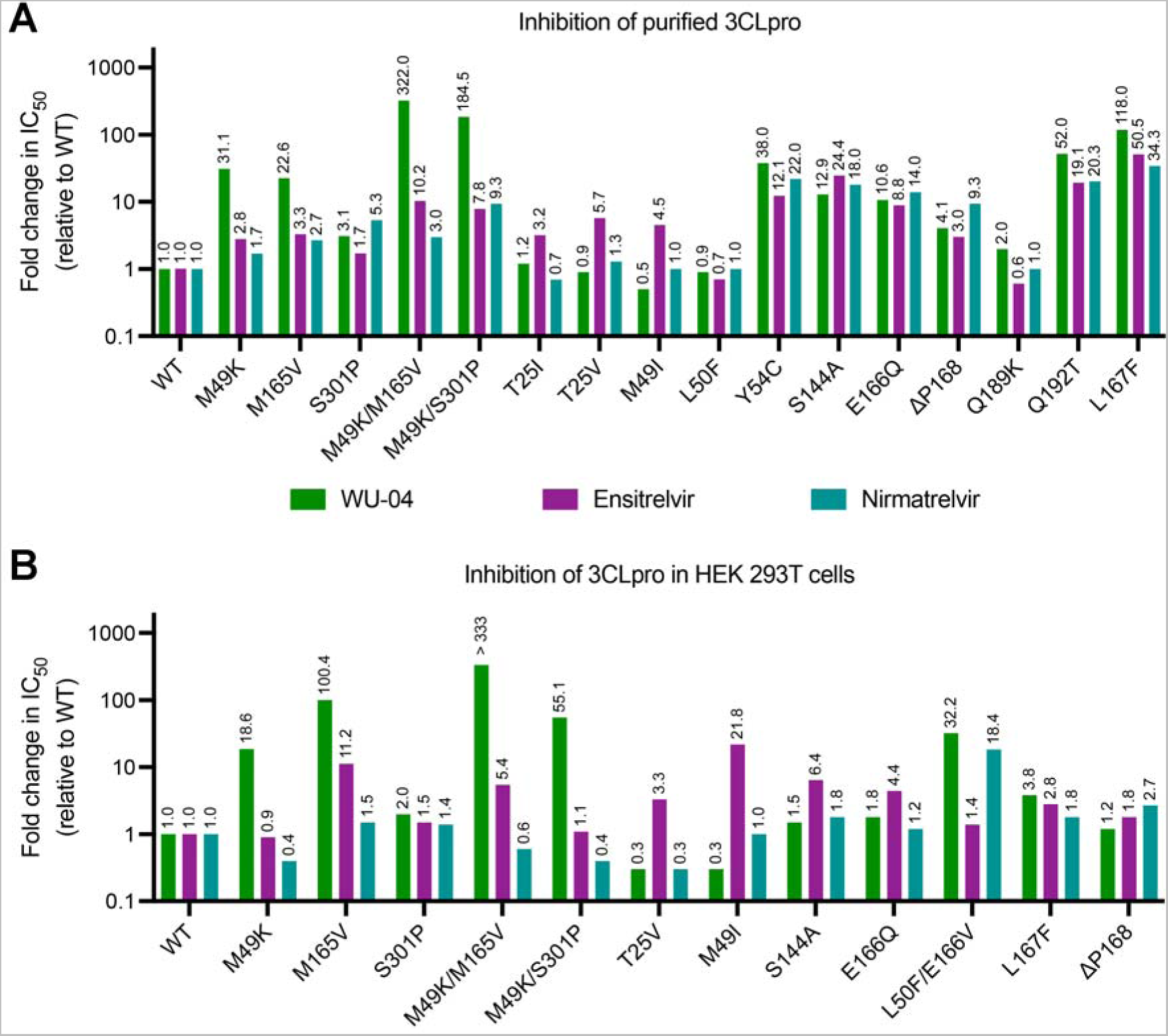
Cross-resistance of 3CLpro mutants to WU-04, ensitrelvir and nirmatrelvir. (**A**, **B**) The inhibitory activities (IC_50_) of WU-04, ensitrelvir and nirmatrelvir against the WT 3CLpro and its mutants were evaluated using a fluorescence-based enzyme assay (**A**), and a BRET-based cell assay (**B**). Each IC_50_ value was calculated based on the data from three independent experiments, then normalized to the IC_50_ value of the corresponding compound against the WT 3CLpro to obtain the fold change value.

For the WU-04-resistant mutants that were identified in our study, they also exhibited differing levels of resistance to ensitrelvir and nirmatrelvir, although the degree of resistance was not as notable as that observed towards WU-04 (Fig. 5A). Specifically, the IC_50_ values of ensitrelvir against the M49K, M165V, and S301P mutants were 2.8, 3.3, and 1.7 times that against the WT 3CLpro, respectively. The double mutants M49K/M165V and M49K/S301P exhibited stronger resistance to ensitrelvir as compared to these single mutants: the IC_50_ values are 10.2 and 7.8 times that against the WT 3CLpro, respectively. An increase in the *K*_i_ of nirmatrelvir was also observed, particularly for the S301P and M49K/S301P mutants.

For the three mutants, including T25I, T25V and M49I, that were selected as ensitrelvir resistant mutants, the IC_50_ values of ensitrelvir against them were 3.2, 5.7, and 4.5 times that against the WT 3CLpro, respectively (Fig. 5A). In contrast, these mutants exhibited negligible resistance to WU-04 and nirmatrelvir. The M49I mutant became even more sensitive to WU-04, with an IC_50_ value approximately half of that against the WT 3CLpro (Fig. S6).

Among the fifteen mutants selected for nirmatrelvir resistance, the catalytic activities of seven were too low to be accurately determined in the enzyme assay. The drug resistance of the remaining eight mutants were evaluated (Fig. 5A). Six mutants, including Y54C, S144A, E166Q, Q192T, L167F, and ΔP168, demonstrated strong resistance to all three inhibitors. The Q189K mutant exhibited moderate resistance to WU-04, but no resistance to ensitrelvir or nirmatrelvir.

The L50F mutant was not resistant to any of the three inhibitors.

### Cross-resistance of 3CLpro mutants expressed in HEK 293T cells to non-covalent and covalent inhibitors

The purified WT 3CLpro and its mutants underwent N- and C-terminal cleavage, thus representing the mature state of 3CLpro. In order to simulate the self-cleavage process of 3CLpro during coronavirus replication, we developed a bioluminescence resonance energy transfer (BRET)-based self-cleaving biosensor pBRET-10, in which a green fluorescent protein (GFP2) and a Renilla luciferase (RLuc8) were linked to the N- and C-termini of SARS-CoV-2 3CLpro, respectively, using linkers derived from the cleavage sequences of 3CLpro (*22*). BRET from RLuc8 to GFP2 can be disrupted by self-cleavage and preserved if the self-cleavage is inhibited. The biosensors carrying the WT 3CLpro and a chosen set of 3CLpro mutants were transiently expressed in human embryonic kidney (HEK) 293T cells. Then the cells were treated with serial dilutions of the inhibitors, and the BRET signals were measured (Fig. S8). The IC_50_ of each inhibitor against each 3CLpro mutant was calculated and normalized to that against the WT 3CLpro (Fig. 5B).

Four mutants that exhibited strong resistance to WU-04 in the enzyme assay, including the M49K, M165V, M49K/M165V, and M49K/S301P mutants, also exhibited strong resistance to WU-04 in the cell assay (Fig. 5B). However, the ranking of resistance levels is different from that in the enzyme assay. Particularly, the M165V mutant demonstrated greater resistance to WU-04 than the M49K and M49K/S301P mutants in the cell assay, whereas its resistance level was lower than that of the M49K and M49K/S301P mutants in the enzyme assay. The M165V mutant also showed greater resistance to ensitrelvir, while the M49K and M49K/S301P mutants showed negligible resistance to ensitrelvir (Fig. 5B). The activity of nirmatrelvir against the four mutants was either moderately affected or even surpassed that against the WT 3CLpro (Fig. 5B).

The T25V and M49I mutants, which exhibited ensitrelvir resistance in the enzyme assay, were also resistant to ensitrelvir in the cell assay, with the respective IC_50_ values 3.3 and 21.8 times that against the WT 3CLpro. Conversely, the two mutants exhibited increased sensitivity to WU-04, with their IC_50_ values decreased to approximately 30% of that against the WT 3CLpro (Fig. 5B); this observation aligns with the findings from the enzyme assay.

Three mutants, S144A, E166Q, and L167F, showed moderate resistance to all the three inhibitors in the cell assay, in contrast to their strong resistance against the three inhibitors in the enzyme assay (Fig. 5B). Among them, S144A and E166Q showed greater resistance to ensitrelvir than to WU-04 and nirmatrelvir. Another mutant, Δ168, exhibited moderate resistance to all three inhibitors (Fig. 5B).

The double mutant L50F/E166V, which showed remarkably weak activity in the enzyme assay, exhibited strong resistance to WU-04 and nirmatrelvir, with the IC_50_ values 32.2 and 18.4 times that against the WT 3CLpro, respectively (Fig. 5B). In contrast, it exhibited negligible resistance to ensitrelvir (Fig. 5B).

## DISCUSSION

We have identified two double mutants of 3CLpro — M49K/M165V and M49K/S301P — that exhibit resistance to the non-covalent inhibitor WU-04. The three mutations in these double mutants each uniquely impact the structure and function of 3CLpro and confer different levels of resistance to WU-04.

Both M49 and M165 are situated within 5 Å of WU-04 (Fig. 1A); mutations at the two residues substantially decrease the binding affinity between 3CLpro and WU-04 (Fig. 1C). The M49K mutation disturbs three regions in 3CLpro (Fig. 2A). Particularly, this mutation destabilizes the short helix of 3CLpro containing residues 45–51, resulting in alterations to the conformation of the WU-04 binding pocket and thereby explaining the resistance to WU-04. In contrast, the crystal structure of the M165V mutant is almost identical to that of the WT 3CLpro (Fig. 2C). M165, located within a β-strand deep within the WU-04 binding pocket, when mutated to the branched-chain amino acid valine, may cause a clash with the bromophenyl ring of WU-04, directly impeding WU-04 binding.

Compared to the M49K and M165V mutations, the S301P mutation had a moderate effect on the binding affinity between 3CLpro and WU-04 (Fig. 1C) and only caused a slight increase in the IC_50_ of WU-04 (Fig. 1B). However, when combined with the M49K mutation, the S301P mutation significantly increased the IC_50_ of WU-04 (Fig. 1B). The S301P mutation restricts the rotation of the C-terminal tail of 3CLpro, thereby perturbing the dimerization of 3CLpro (Fig. 3). This unique resistance mechanism is different from those of the M49K and M165V mutations. The S301P mutation was also identified in the screening for nirmatrelvir-resistant mutations (*8*). We confirmed that it conferred resistance to nirmatrelvir (Fig. 5A). This observation indicates that restriction of the rotation of the C-terminal tail of 3CLpro serves as a resistance mechanism for nirmatrelvir as well.

Alongside the WU-04-resistant mutations, we also investigated mutations and deletions that confer resistance to ensitrelvir and nirmatrelvir. Almost all of these alterations resulted in a decrease in the catalytic activity of 3CLpro (Fig. 4G). Specific mutations, including E166V, L50F/E166V, H172Y, H163W, M165Y, ΔQ189, and ΔQ192, significantly reduced the catalytic activity of 3CLpro to almost undetectable levels (Fig. 4G and Fig. S4). Most of these mutations also decreased the thermostability of 3CLpro (Fig. 4H). These findings suggest a trade-off between the catalytic activity, thermostability and drug resistance of 3CLpro. However, even with low levels of 3CLpro protease activity, viral replication can still be sustained, as evidenced by the high fitness of the virus carrying either the L50F/E166V double mutation or the L50F/E166A/L167F triple mutation in 3CLpro (*9, 23*), suggesting that nearly complete inhibition is needed to block viral replication.

An unexpected finding is that the L50F mutation, previously reported to significantly decrease the catalytic activity of 3CLpro (*7, 9*), actually enhanced the catalytic activity of 3CLpro (Fig. 4G and Fig. S4). Our finding aligns with the observation that the L50F mutation compensated for the replicative fitness loss caused by the E166V mutation (*8, 9*). A yeast screen also indicated that the L50F mutation can increase the catalytic activity of 3CLpro, however, there was a lack of enzymatic assay data (*17*). A recent study showed that the double mutant L50F/E166V had increased catalytic activity compared to the E166V single mutant, indicating that the L50F mutation increased the 3CLpro catalytic activity (*18*). The controversy over the catalytic activity of the L50F mutant may be due to the difficulty in obtaining a well-behaved, purified sample of this mutant in previous studies. We successfully expressed and purified the L50F mutant, thus presenting conclusive evidence of its increased catalytic activity. This finding suggests that the decreased catalytic activity of 3CLpro caused by most of the drug-resistant mutations can be restored by introducing additional mutations.

A number of mutations, such as Y54C, S144A, E166Q, Q192T, and L167F, confer robust resistance to all three inhibitors. However, certain mutants exhibit resistance to one inhibitor while remaining sensitive to others, as exemplified by mutations at M49 (Fig. 5). In enzyme assays, the M49K mutation significantly increased the IC_50_ of WU-04, but only moderately affected the potency of ensitrelvir and nirmatrelvir; conversely, the M49I mutation had minimal impact on the potency of WU-04 and nirmatrelvir but caused a significant increase in the IC_50_ of ensitrelvir (Fig. 5A). In our cell-based assay that was designed to evaluate the self-cleavage efficiency of 3CLpro, the M49K mutation resulted in a nearly 20-fold increase in the IC_50_ of WU-04 but had negligible effect on the efficacy of ensitrelvir (Fig. 5B). In contrast, the M49I mutation caused a nearly 20-fold increase in the IC_50_ of ensitrelvir but decreased the IC_50_ of WU- 04 (Fig. 5B). These findings demonstrate that mutations at the same residue can have distinct effects on the 3CLpro inhibitors. This underscores the importance of developing multiple antiviral agents with different skeletons for fighting SARS-CoV-2.

We used two assays to evaluate the resistance conferred by 3CLpro mutations: an enzyme assay using purified, mature 3CLpro, and a cell assay using a biosensor to mimic the maturation process of 3CLpro in HEK 293T cells. Interestingly, these two assays yielded different rankings of resistance levels for the 3CLpro mutations. For instance, while the L167F mutation conferred stronger resistance to WU-04 than the M49K and M165V mutations in the enzyme assay, it demonstrated much weaker resistance in the cell assay. This discrepancy potentially suggests that the inhibitory activity varies between mature and unprocessed 3CLpro. However, the precise molecular mechanisms underpinning this difference remain to be elucidated.

## ACKNOWLEDGMENTS

We thank Dr. Sheng-ce Tao at Shanghai Jiao Tong University for sharing the plasmids of SARS-CoV-2 proteins. We thank the Protein Characterization and Crystallography Facility of Westlake University for assistance in crystallization and X-ray data collection. We thank Shan Feng and Jinheng Pan at the Mass Spectrometry & Metabolomics Core Facility of Westlake University for their technical assistance. We would like to express our sincere gratitude towards Tencent Foundation for their generous support in this collaborative project. This work was supported by Westlake Laboratory of Life Sciences and Biomedicine (to J.H., Q.H.), Central Guidance on Local Science and Technology Development Fund (2022ZY1006, to Q.H.), Westlake Education Foundation (to J.H., Q.H.), Tencent Foundation (to J.H., Q.H.).

## AUTHOR CONTRIBUTIONS

Q.H. conceived the project. L.Z. designed and performed the biochemical assays. L.Z. and H.L. purified the proteins and did the crystallization experiments. Q.H. and L.Z. determined and refined the crystal structures. X.X. and P.Y.S. performed virology experiments. All the authors analyzed the data. L.Z. and Q.H. wrote the manuscript.

## COMPETING INTEREST STATEMENT

L.Z., H.Y., J.H., and Q.H. are co-inventors of patents that cover the 3CLpro inhibitor WU-04 in this study. H.Y., J.H., and Q.H. are founders of Westlake Pharmaceuticals (Hangzhou) Co., Ltd. and members of its scientific advisory board. Other authors declare no competing interests.

## REFERENCES

1. A. R. Fehr, S. Perlman, Coronaviruses: an overview of their replication and pathogenesis. Methods Mol Biol 1282, 1–23 (2015).

2. P. Zhou et al., A pneumonia outbreak associated with a new coronavirus of probable bat origin. Nature 579, 270–273 (2020).

3. V. Coronaviridae Study Group of the International Committee on Taxonomy of, The species Severe acute respiratory syndrome-related coronavirus: classifying 2019-nCoV and naming it SARS-CoV-2. Nat Microbiol 5, 536–544 (2020).

4. D. R. Owen et al., An oral SARS-CoV-2 M(pro) inhibitor clinical candidate for the treatment of COVID-19. Science 374, 1586–1593 (2021).

5. Y. Unoh et al., Discovery of S-217622, a Noncovalent Oral SARS-CoV-2 3CL Protease Inhibitor Clinical Candidate for Treating COVID-19. J Med Chem 65, 6499–6512 (2022).

6. J. Ou et al., A yeast-based system to study SARS-CoV-2 Mpro structure and to identify nirmatrelvir resistant mutations. Res Sq, (2022).

7. D. Jochmans et al., The Substitutions L50F, E166A, and L167F in SARS-CoV-2 3CLpro Are Selected by a Protease Inhibitor In Vitro and Confer Resistance To Nirmatrelvir. mBio 14, e0281522 (2023).

8. S. Iketani et al., Multiple pathways for SARS-CoV-2 resistance to nirmatrelvir. Nature 613, 558–564 (2023).

9. Y. Zhou et al., Nirmatrelvir-resistant SARS-CoV-2 variants with high fitness in an infectious cell culture system. Sci Adv 8, eadd7197 (2022).

10. Y. Hu et al., Naturally occurring mutations of SARS-CoV-2 main protease confer drug resistance to nirmatrelvir. bioRxiv, (2022).

11. V. M. Sasi et al., Predicting Antiviral Resistance Mutations in SARS-CoV-2 Main Protease with Computational and Experimental Screening. Biochemistry 61, 2495–2505 (2022).

12. K. S. Yang, S. Z. Leeuwon, S. Xu, W. R. Liu, Evolutionary and Structural Insights about Potential SARS-CoV-2 Evasion of Nirmatrelvir. J Med Chem 65, 8686–8698 (2022).

13. S. Iketani et al., Functional map of SARS-CoV-2 3CL protease reveals tolerant and immutable sites. Cell Host Microbe 30, 1354–1362 e1356 (2022).

14. E. Heilmann et al., SARS-CoV-2 3CL(pro) mutations selected in a VSV-based system confer resistance to nirmatrelvir, ensitrelvir, and GC376. Sci Transl Med 15, eabq7360 (2023).

15. S. A. Moghadasi et al., Transmissible SARS-CoV-2 variants with resistance to clinical protease inhibitors. Sci Adv 9, eade8778 (2023).

16. G. D. Noske et al., Structural basis of nirmatrelvir and ensitrelvir activity against naturally occurring polymorphisms of the SARS-CoV-2 main protease. J Biol Chem 299, 103004 (2023).

17. J. M. Flynn et al., Systematic Analyses of the Resistance Potential of Drugs Targeting SARS-CoV-2 Main Protease. ACS Infect Dis 9, 1372–1386 (2023).

18. Y. Duan et al., Molecular mechanisms of SARS-CoV-2 resistance to nirmatrelvir. Nature 622, 376–382 (2023).

19. N. Hou et al., Development of Highly Potent Noncovalent Inhibitors of SARS-CoV-2 3CLpro. ACS Cent Sci 9, 217–227 (2023).

20. M. Kiso et al., In vitro and in vivo characterization of SARS-CoV-2 resistance to ensitrelvir. Nat Commun 14, 4231 (2023).

21. F. H. Niesen, H. Berglund, M. Vedadi, The use of differential scanning fluorimetry to detect ligand interactions that promote protein stability. Nat Protoc 2, 2212–2221 (2007).

22. N. Hou, C. Peng, L. Zhang, Y. Zhu, Q. Hu, BRET-Based Self-Cleaving Biosensors for SARS-CoV-2 3CLpro Inhibitor Discovery. Microbiol Spectr 10, e0255921 (2022).

23. R. Abdelnabi et al., Nirmatrelvir-resistant SARS-CoV-2 is efficiently transmitted in female Syrian hamsters and retains partial susceptibility to treatment. Nat Commun 14, 2124 (2023).

24. B. A. Johnson et al., Nucleocapsid mutations in SARS-CoV-2 augment replication and pathogenesis. PLoS Pathog 18, e1010627 (2022).

25. X. Xie et al., An Infectious cDNA Clone of SARS-CoV-2. Cell Host Microbe 27, 841–848 e843 (2020).

