## Supplementary material for "Resistance mechanisms of SARS-CoV-2 3CLpro to the non-covalent inhibitor WU-04": Materials and Methods, Supplemental Figures S1 to S8, Supplemental Tables S1 and S2.

**Affiliations:**

### MATERIALS AND METHODS

#### Selection and sequencing of WU-04 resistant virus

The WU-04-resistant virus was obtained by serial passaging of the SARS-CoV-2 mNG (this virus showed significantly reduced virulence in vivo compared to the wild-type SARS-CoV-2) (*24*) in Vero E6 cells in the presence of increasing concentrations of WU-04. Four selections for WU-04-resistant virus were independently performed. Briefly, a reporter virus, named SARS-CoV-2 mNG, was generated by introducing the gene of mNeonGreen into ORF7 of the SARS-CoV-2 viral genome (*25*). For each selection, Vero E6 cell monolayers in a 12-well plate were inoculated at a multiplicity of infection (MOI) of 2 with SARS-CoV-2 mNG or previously passaged virus and the compound. After incubation at 37 °C for 1 hour, the inoculum was removed, and 1 mL fresh DMEM medium with 2% FBS containing WU-04 was added to each well. The cell cultures were incubated at 37 °C for 2-5 days. The culture medium was harvested when over 80% of cells showed mNG-positive. Selection began at the WU-04 concentration of 120 nM (P1) and was followed by increasing the WU-04 concentrations to 480 nM (P2), 2000 nM (P3), 4000 nM (P4), and 10000 nM (P5-P7). The supernatant from 10000 nM WU-04-selected virus (P7) was tested for WU-04 sensitivity on Vero CCL81 cells. For viral genome sequencing, viral RNA was extracted from P7 culture fluids by using the TRIzol™ LS Reagent (ThermoFisher Scientific), and the cDNA fragment containing the nucleotides 7382-11990 in the SARS-CoV-2 genome was amplified by using SuperScript™ IV One-Step RT-PCR System (ThermoFisher Scientific). The RT-PCR products were purified and subjected to Sanger sequencing. The selection process for the WU-04 resistant viruses was conducted during the early stage of COVID-19 pandemic, before any antiviral drugs, including nirmatrelvir, had been authorized for emergency use. During the selection process, WU-04 resistant viruses exhibited comparable replication kinetics to the SARS-CoV-2 mNG strain. The selected viruses had only been tested for their sensitivity to WU-04. No other antiviral inhibitors have been tested using these resistant viruses. All virus experiments were conducted at Biosafety level-3 laboratories. Personnel who performed the experiments wore powered air-purifying respirators (Breathe Easy, 3M) with Tyvek suits, aprons, booties, and double gloves.

**Genes and cloning**

The gene coding the SARS-CoV-2 3CLpro was a gift from Prof. Sheng-ce Tao at Shanghai Jiao Tong University. The plasmid of SARS-CoV-2 3CLpro (wild type) for protein expression and BRET assay were reported in our previous studies (*19, 22*). The plasmids of SARS-CoV-2 3CLpro mutants were constructed through QuickChange site-directed mutagenesis by using ClonExpress® II One Step Cloning Kit (Vazyme, C112-02).

**Protein expression and purification**

The SARS-CoV-2 3CLpro proteins were overexpressed in *Escherichia coli* and purified following a method described previously (*19*). In brief, the His-tagged proteins were purified by affinity chromatography using $Co^{2+}$ resin (TALON, Cat# 635504), then the tag was removed by human rhinovirus 3C protease (TaKaRa, Cat# 7360) and the efficiency of cleavage was analyzed by SDS-PAGE (GenScript, M00656). The proteins were further purified by ion-exchange chromatography (Source-15Q column, GE Healthcare) and size-exclusion chromatography (Superdex 200 increase 10/300 GL column, GE Healthcare). Finally, the purified proteins were concentrated and stored in 20 mM HEPES, pH 7.4, 150 mM NaCl at -80 °C for subsequent biological assays and crystallization.

**3CLpro FRET-based assay**

The enzymatic activities of the WT 3CLpro and its mutants, and the inhibitory activity of each inhibitor (WU-04, Ensitrelvir, and Nirmatrelvir) were evaluated using a fluorescence resonance energy transfer (FRET)-based assay as described previously (*19*). The fluorogenic peptide Dabcyl-KTSAVLQSGFRKME-Edans was used as the substrate. For the enzyme kinetic study, the final concentration of 3CLpro was 25 nM. In detail, 20 μL of 3CLpro (50 nM) in the reaction buffer (20 mM HEPES 7.4, 150 mM NaCl, 0.01% Triton X-100, 1 mM DTT) was added into a 384-well black plate (Corning, CLS3575) and incubated at 37 °C for 5 minutes, then 20 μL of different concentrations of fluorogenic substrate in the reaction buffer (0-200 μM) was added to each well to initiate the reaction. The fluorescence was monitored at 37 °C using an excitation wavelength of 355 nm and an emission wavelength of 538 nm in a Thermol Varioskan LUX plate reader. A control experiment containing only the fluorogenic substrate in the reaction was carried out. A standard curve was generated using the product (SGFRKME-Edans), then the fluorescence signals of each sample were converted to the product concentrations. The slope of each curve from 0 to 10 minutes was calculated as the velocity of the corresponding reaction. Three independent experiments were performed. The data were fitted by the software GraphPad Prism 9 using the Michaelis-Menten equation. To measure the $IC_{50}$ values of WU-04 and Ensitrelvir, the final concentration of 3CLpro was 25 nM. Specially, 10 μL of each inhibitor at a series of concentrations in the dilution buffer (20 mM HEPES 7.4, 150 mM NaCl, 0.01% Triton X-100, 1 mM DTT, 10% DMSO) was incubated with 10 μL of 3CLpro (100 nM) in the reaction buffer (20 mM HEPES 7.4, 150 mM NaCl, 0.01% Triton X-100, 1 mM DTT) at room temperature for 30 min and then incubated at 37 °C for 5 minutes. Next, 20 μL of the fluorogenic substrate (50 μM) in the reaction buffer was added to each well to initiate the reaction. The fluorescence was monitored at 37 °C with an excitation wavelength of 355 nm and an emission wavelength of 538 nm using a Thermol Varioskan LUX plate reader. A control experiment containing inhibitors and the fluorogenic substrate in the reaction was carried out. The slope of each fluorescence curve from 0 to 10 minutes was calculated as the velocity of the corresponding reaction. Three independent experiments were performed. The data was analyzed using a four-parameters model in GraphPad Prism 9 software. To measure the $K_i$ values of Nirmatrelvir, 10 µL of Nirmatrelvir at a series of concentrations in the dilution buffer (20 mM HEPES 7.4, 150 mM NaCl, 0.01% Triton X-100, 1 mM DTT, 10% DMSO) was incubated with 20 µL of the fluorogenic substrate (50 µM) in the reaction buffer at 37 °C for 5 minutes. Then 10 µL of 3CLpro (100 nM) in the reaction buffer (20 mM HEPES 7.4, 150 mM NaCl, 0.01% Triton X-100, 1 mM DTT) was added to each well to initiate the proteolytic reaction. The $K_i$ was calculated by plotting the initial velocity against the concentration of Nirmatrelvir using the Morrison $K_i$ plot in Prism 9 software.

**Isothermal titration calorimetry (ITC)**

ITC experiments were done with the isothermal titration calorimeter MicroCal PEAQ-ITC (Malvern Panalytical). 20 µM of WU-04 in the ITC buffer (20 mM HEPES, pH 7.4, 150 mM NaCl, 0.5% DMSO) was titrated by 200 µM of 3CLpros in the ITC buffer at 25 °C. The data was processed using the MicroCal PEAQ-ITC analysis software.

**Crystallization**

The SARS-CoV-2 3CLpro mutants were concentrated to 10 mg/ml, followed by centrifugation at 21,000 g for 5 minutes to remove the precipitate. DTT was added to a final concentration of 5 mM before crystallization for M49K/S301P. For crystallization, 0.2 µL of the protein was mixed with 0.2 µL of well buffer in a 96-well plate by a protein crystallization robot (Mosquito) using the sitting drop method (M165V and S301P) or hanging drop method (M49K, M49K/165V and M49K/S301P), then the drop was equilibrated against 90 µL of the well buffer at 20 °C. The well buffer for the crystallization of the M49K mutant contained 0.2 M BIS-TRIS, pH 6.0, 20% w/v polyethylene glycol 4,000. The well buffer for the crystallization of the M165V mutant contained 0.2 M BICINE, pH 8.1, 20% polyethylene glycol 4,000. The well buffer for the crystallization of the S301P mutant contained 0.2 M BIS-TRIS, pH 6.6, 20% polyethylene glycol 4,000. The well buffer for the crystallization of the M49K/M165V double mutant contained 0.2 M BIS-TRIS propane, pH 7.3, 20% polyethylene glycol 4,000. The well buffer for the crystallization of the M49K/S301P double mutant contained 0.2 M LiSO4, 0.1 M BIS-TRIS, pH 6.6, 17.5% polyethylene glycol 3,350. The complex of the M49K/S301P double mutant with WU-04 was prepared by incubating the M49K/S301P double mutant (10 mg/ml in 20 mM HEPES, pH7.4, 150 mM NaCl) with 1.5 mM WU-04 (the stock used is 50 mM in DMSO) at room temperature for 2 hours, followed by centrifugation at 21,000 g for 5 minutes to remove the precipitate. Then, 0.2 µL of the complex was mixed with 0.2 µL of the well buffer in a 96-well plate using the sitting drop method and the drop was equilibrated against 90 µL of the well buffer at 20 °C. The well buffer contains 0.1 M sodium formate, 12% polyethylene glycol 3,350.

**Data collection and structure determination**

The crystals were first transferred to a cryoprotectant solution (the well buffer plus 20 mM HEPES, pH 7.4, 150 mM NaCl, and 10-20% glycerol), then loaded onto the X-ray diffractometer (Rigaku, XtaLAB Synergy Customer) at Westlake University. The diffraction data was collected at 100 K and processed with the reduction program CrysAlisPro. The structures were solved by molecular replacement using PHaser in PHENIX. The co-crystal structure of SARS-CoV-2 3CLpro/WU-04 (PDB ID code 7EN8) was used as the initial model. The structures were manually refined with Coot and PHENIX. Data collection and refinement statistics can be found in table S1 that was generated using the utility PHenix.table_one in PHENIX.

**Bioluminescence resonance energy transfer (BRET) based cell assay**

The BRET assay was performed following a method described previously (*22*). Briefly, HEK 293T cells were seeded into a 96-well clear-bottom white plate (Corning, 3903) at approximately 40% confluence. After 24 hours, the cells were transfected with plasmids carrying the biosensors (400 ng/well) using PEI as the transfection reagent. Then, 3CLpro inhibitors with a series of concentrations in DMSO were added to the cell culture. The well with the HEK 293T cells transfected with an empty plasmid and treated with DMSO was used as a blank control. The final concentration of DMSO in the cell culture was 0.48%. 24 hours post transfection, coelenterazine 400a (GoldBio, C-320) was added to reach a final concentration of 10 μM and the luminescence at 413 nm (wavelength range from 400 nm to 425 nm) and fluorescence at 518 nm (wavelength range from 505 nm to 530 nm) were measured after shaking for 5 s using a plate reader (TECAN-Spark). The BRET ratio was calculated using the following equation:

$$\text{BRET ratio} = (F_{518,\,S} - F_{518,\,BLK})/(L_{413,\,S} - L_{413,\,BLK}),$$

in which $F_{518,\,S}$ and $L_{413,\,S}$ are the fluorescence (518 nm) and luminescence (413 nm) signals cells, respectively, while $F_{518,\,BLK}$ and $L_{413,\,BLK}$ are the fluorescence (518 nm) and luminescence (413 nm) signals of the blank control.

**Thermal shift assay**

The thermal shift assay was performed in the BIO-RAD CFX Connect Real-Time PCR Detection System. To 10 μL of 3CLpro (10 μM) in 20 mM HEPES, pH 7.4, 150 mM NaCl in a 96-well PCR plate (BIO-RAD MLL9601), 10 μL of 10x SYPRO™ Orange in the reaction buffer (20 mM HEPES, pH 7.4, 150 mM NaCl, 2 mM DTT, 0.02% Triton X-100) was added and mixed gently. The final reaction contained 5 μM of 3CLpro, 5x SYPRO™ Orange, 20 mM HEPES, pH 7.4, 150 mM NaCl, 1 mM DTT, 0.01% Triton X-100. The reaction buffer plus SYPRO™ Orange was used as no protein control. The fluorescence was monitored under a temperature gradient ranging from 25 to 95 °C in 0.5 °C increments every 30 s after an initial incubation at 25 °C for 5 min. Each data set was normalized to the highest fluorescence, and the normalized fluorescence reading was plotted against temperature in GraphPad Prism 9. The melting temperature ($T_m$) values were determined as the temperature corresponding to the maximum of the first derivative of the curve. The melting temperature shift ($\Delta T_m$) of 3CLpro mutants was calculated by subtracting the $T_m$ of the wild type 3CLpro.

**A**

WU-04 resistant mutations in 3CLpro

| Viruses | $EC_{50}$s (nM) | 3CL mutations |
|---|---|---|
| WT mNG | 18.2, 17.2 | NA |
| 14h6 P7-1 | > 5000 | T10200A (3CL: M49K), A10547G (3CL: M165V) |
| 14h6 P7-2 | > 5000 | T10200A (3CL: M49K), A10547G (3CL: M165V) |
| 14h6 P7-3 | > 5000 | T10200A (3CL: M49K), T10955C (3CL: S301P) |
| 14h6 P7-4 | > 5000 | T10200A (3CL: M49K), T10955C (3CL: S301P) |

**B**

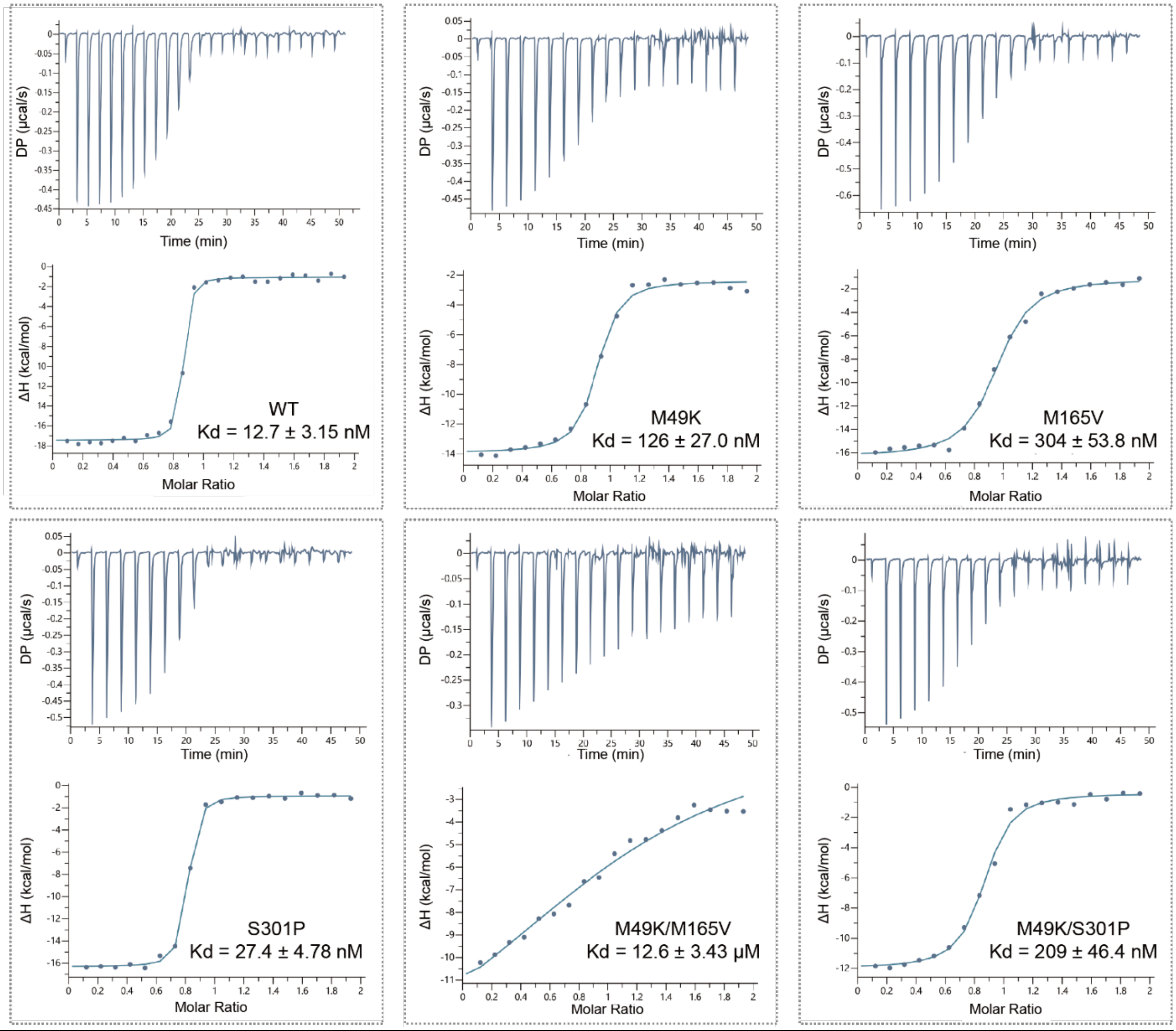


**Fig. S1.** Identification of WU-04-resistant mutations in SARS-CoV-2 3CLpro. (**A**) The WU-04-resistant viruses were identified by serial passaging of the SARS-CoV-2 mNG in Vero E6 cells treated with increasing concentrations of WU-04. (**B**) The binding affinities ($K_d$) between WU-04 and the WU-04-resistant 3CLpro mutants were measured using isothermal titration calorimetry (ITC).

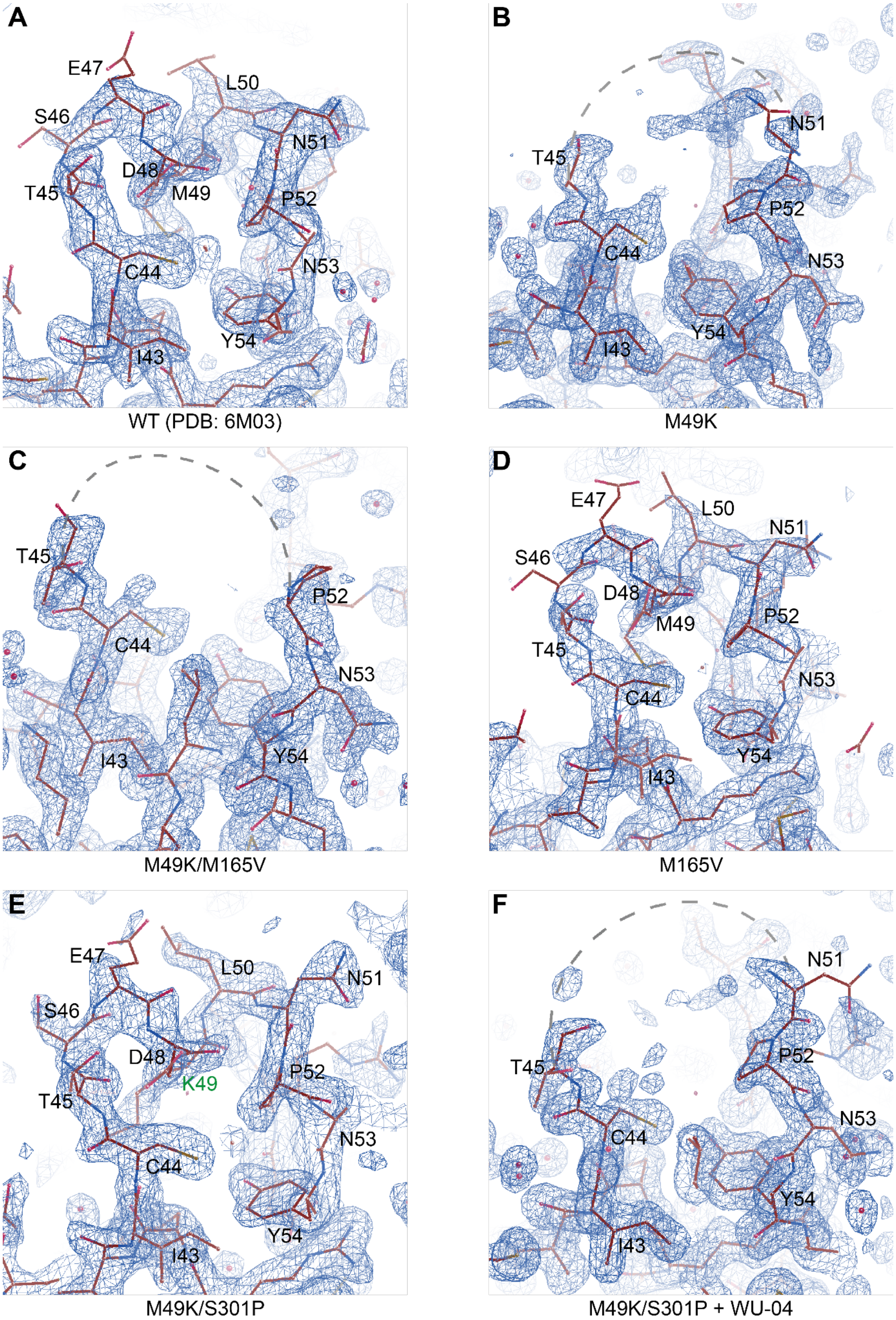
A
E47
L50
S46
N51
D48
M49
T45
P52
C44
N53
Y54
I43
WT (PDB: 6M03)
B
N51
T45
P52
C44
N53
Y54
I43
M49K
C
T45
P52
C44
N53
I43
Y54
M49K/M165V
D
E47
L50
N51
S46
D48
M49
P52
T45
N53
C44
Y54
I43
M165V
E
E47
L50
N51
S46
D48
K49
P52
T45
N53
C44
Y54
I43
M49K/S301P
F
N51
T45
P52
N53
C44
Y54
I43
M49K/S301P + WU-04

**Fig. S2.** The 2mFo-DFc electron density maps of the helix (residues 45-51) of the crystal structures. The *2mFo-DFc* electron density maps of the helix (residues 45-51) of the wild-type SARS-CoV-2 3CLpro (PDB ID code 6M03) (**A**), the WU-04-resistant mutants M49K (**B**), M49K/M165V (**C**), M165V (**D**), M49K/S301P (**E**), and the M49K/S301P mutant in complex with WU-04 (**F**) were contoured at a level of 1.2 RMSD in *coot*.

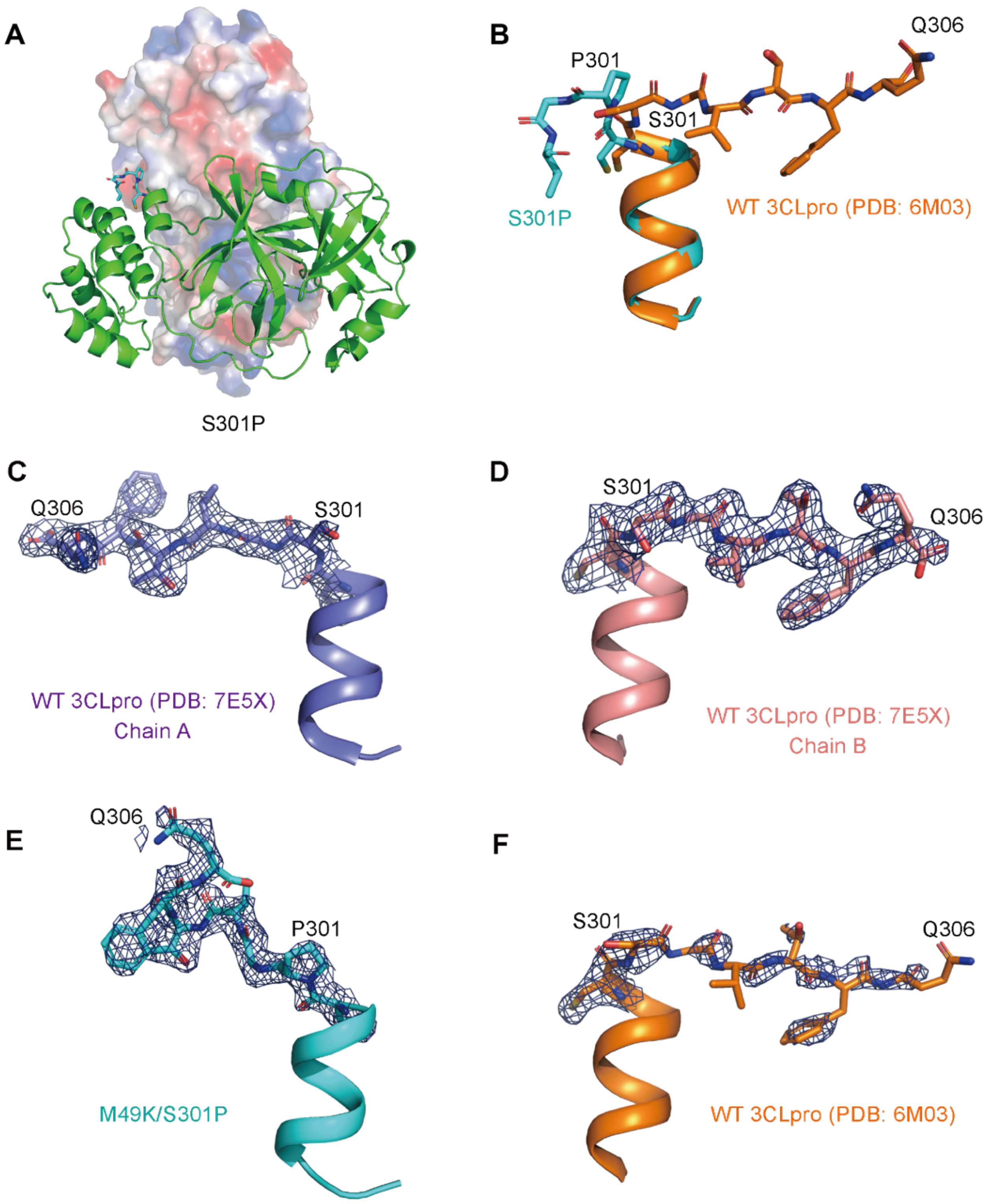


**Fig. S3.** The C-terminal tails (residues 301-306) of 3CLpro in the crystal structures. (**A**) In the crystal structure of the S301P mutant, the C-terminal tail (colored cyan) of one 3CLpro protomer turns away from the other 3CLpro protomer within the same 3CLpro homodimer. (**B**) Alignment of the C-terminal tails of 3CLpro in the S301P structure (colored cyan) with that in the mature WT 3CLpro structure (colored orange). (**C-F**) The *2Fo-Fc* maps of the C-terminal tails (residues 301-306) of the WT 3CLpro in the post-cleavage state (PDB ID code 7E5X) (**C**, **D**), in the crystal structure of the WU-04 04-resistant mutant M49K/S301P (**E**), and that in the crystal structure of the WT 3CLpro in the mature state (PDB ID code 6M03) (**F**) were made using PyMOL. The contour level was 1.0 σ.

A

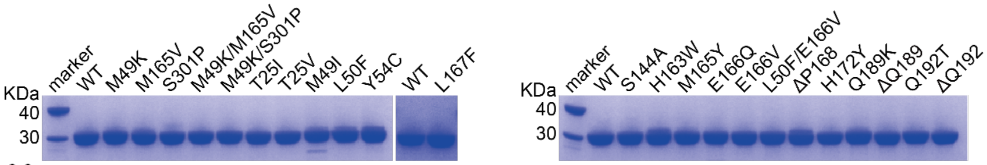


B

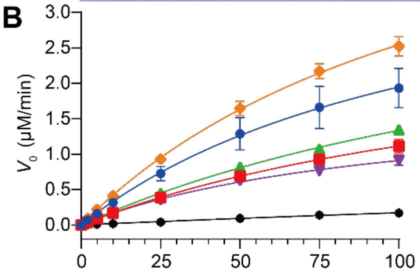


| | $V_{max}$ (μM·min$^{-1}$) | $K_m$ (μM) | $k_{cat}$ (min$^{-1}$) | $k_{cat}/K_m$ (μM$^{-1}$·min$^{-1}$) |
|---|---|---|---|---|
| WT | 4.26 ± 0.83 | 119 ± 37 | 170.4 | 1.43 |
| T25I | 2.94 ± 0.33 | 163 ± 27 | 117.6 | 0.72 |
| T25V | 3.87 ± 0.27 | 190 ± 18 | 154.8 | 0.81 |
| M49I | 1.81 ± 0.13 | 100 ± 12 | 72.4 | 0.73 |
| L50F | 5.78 ± 0.41 | 127 ± 14 | 231.2 | 1.82 |
| Y54C | 1.29 ± 0.83 | 651 ± 469 | 51.6 | 0.08 |

C

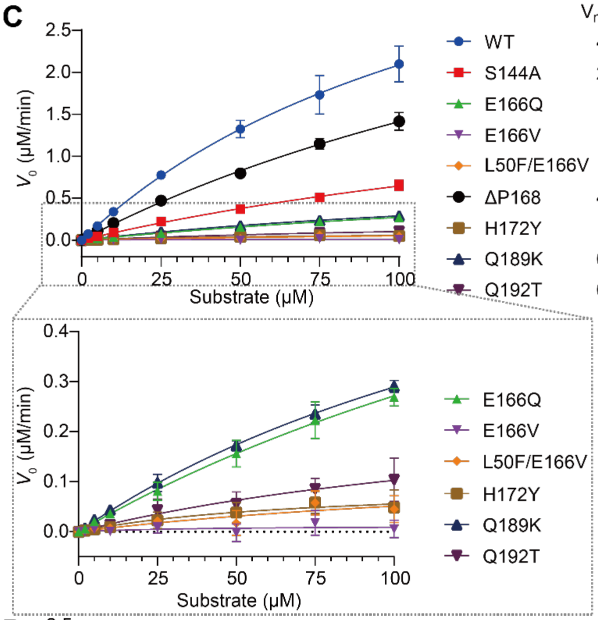


| | $V_{max}$ (μM·min$^{-1}$) | $K_m$ (μM) | $k_{cat}$ (min$^{-1}$) | $k_{cat}/K_m$ (μM$^{-1}$·min$^{-1}$) |
|---|---|---|---|---|
| WT | 4.94 ± 0.71 | 137 ± 30 | 197.6 | 1.44 |
| S144A | 2.25 ± 0.40 | 249 ± 58 | 90 | 0.36 |
| E166Q | 1.03 ± 0.36 | 281 ± 123 | 41.2 | 0.15 |
| E166V | ND | ND | ND | ND |
| L50F/E166V | ND | ND | ND | ND |
| ΔP168 | 4.83 ± 0.78 | 242 ± 52 | 193.2 | 0.8 |
| H172Y | ND | ND | ND | ND |
| Q189K | 0.86 ± 0.12 | 196 ± 38 | 34.24 | 0.17 |
| Q192T | 0.28 ± 0.18 | 172 ± 160 | 11.16 | 0.06 |

D

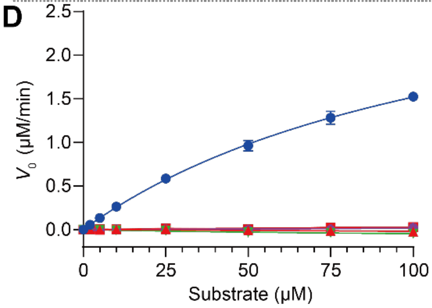


| | $V_{max}$ (μM·min$^{-1}$) | $K_m$ (μM) | $k_{cat}$ (min$^{-1}$) | $k_{cat}/K_m$ (μM$^{-1}$·min$^{-1}$) |
|---|---|---|---|---|
| WT | 3.38 ± 0.21 | 122 ± 12 | 135.2 | 1.11 |
| H163W | ND | ND | ND | ND |
| M165Y | ND | ND | ND | ND |
| ΔQ189 | ND | ND | ND | ND |
| ΔQ192 | ND | ND | ND | ND |

E

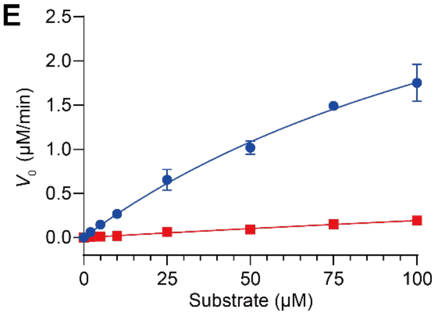


| | $V_{max}$ (μM·min$^{-1}$) | $K_m$ (μM) | $k_{cat}$ (min$^{-1}$) | $k_{cat}/K_m$ (μM$^{-1}$·min$^{-1}$) |
|---|---|---|---|---|
| WT | 4.66 ± 0.78 | 165 ± 40 | 186.4 | 1.13 |
| L167F | 1.89 ± 2.93 | 886 ± 1499 | 75.6 | 0.09 |

**Fig. S4.** The enzymatic activities of SARS-CoV-2 3CLpro mutants. The enzymatic activity of each mutant was evaluated using a FRET-based assay. The data of WT 3CLpro used in Fig. S4B was also used in Fig. 1D. The data represent the mean ± SD of three independent measurements.

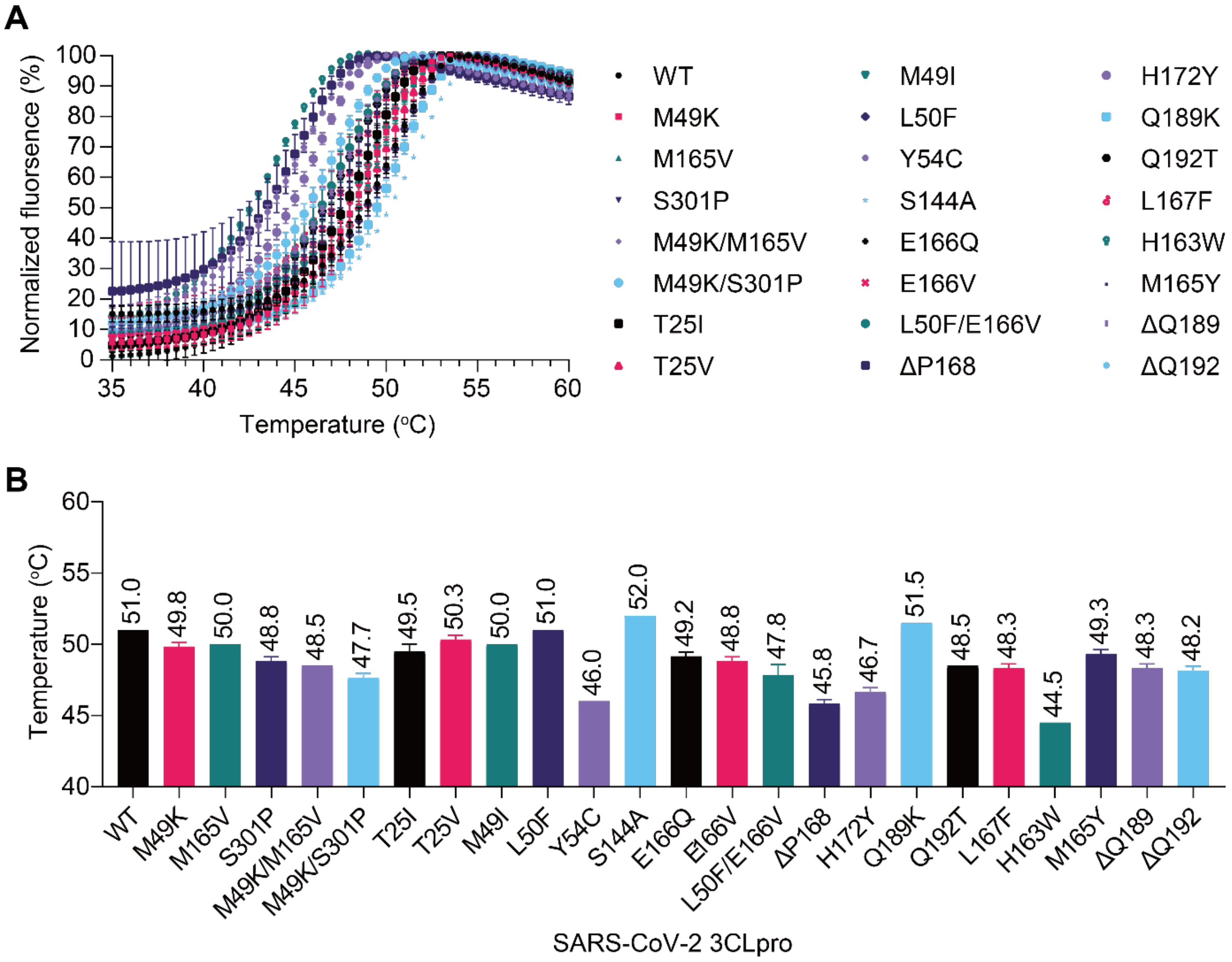


**Fig. S5.** Thermal shift results of the SARS-CoV-2 3CLpro mutants. The data represent the mean ± SD of the technical triplicate. (**A**) The melting curves of the WT 3CLpro and the drug-resistant mutants. (**B**) The calculated melting temperatures of the WT 3CLpro and the drug-resistant mutants

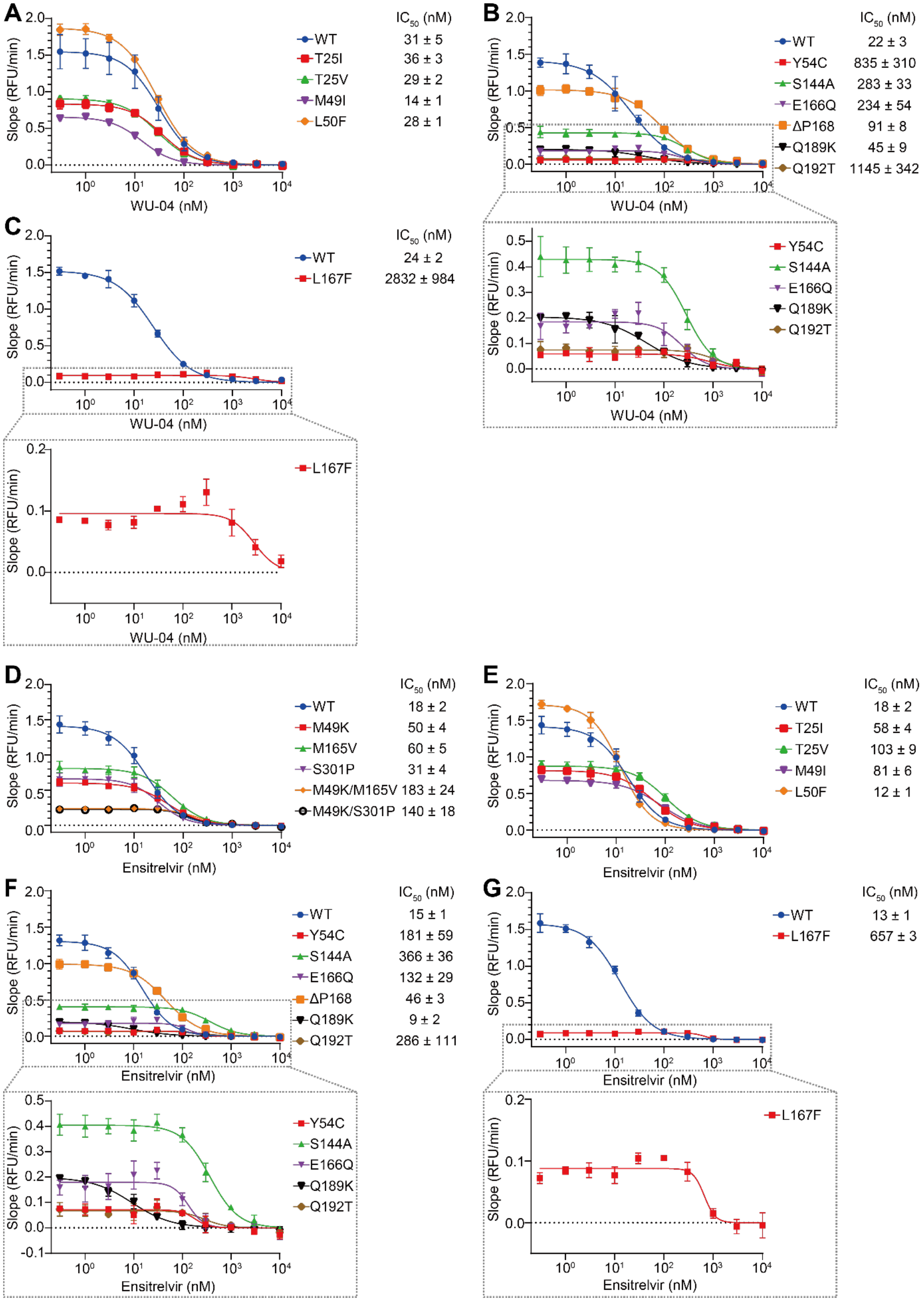
A
Slope (RFU/min)
WU-04 (nM)
IC50 (nM)
WT 31 ± 5
T25I 36 ± 3
T25V 29 ± 2
M49I 14 ± 1
L50F 28 ± 1
B
Slope (RFU/min)
WU-04 (nM)
IC50 (nM)
WT 22 ± 3
Y54C 835 ± 310
S144A 283 ± 33
E166Q 234 ± 54
ΔP168 91 ± 8
Q189K 45 ± 9
Q192T 1145 ± 342
Y54C
S144A
E166Q
Q189K
Q192T
C
Slope (RFU/min)
WU-04 (nM)
IC50 (nM)
WT 24 ± 2
L167F 2832 ± 984
L167F
D
Slope (RFU/min)
Ensitrelvir (nM)
IC50 (nM)
WT 18 ± 2
M49K 50 ± 4
M165V 60 ± 5
S301P 31 ± 4
M49K/M165V 183 ± 24
M49K/S301P 140 ± 18
E
Slope (RFU/min)
Ensitrelvir (nM)
IC50 (nM)
WT 18 ± 2
T25I 58 ± 4
T25V 103 ± 9
M49I 81 ± 6
L50F 12 ± 1
F
Slope (RFU/min)
Ensitrelvir (nM)
IC50 (nM)
WT 15 ± 1
Y54C 181 ± 59
S144A 366 ± 36
E166Q 132 ± 29
ΔP168 46 ± 3
Q189K 9 ± 2
Q192T 286 ± 111
Y54C
S144A
E166Q
Q189K
Q192T
G
Slope (RFU/min)
Ensitrelvir (nM)
IC50 (nM)
WT 13 ± 1
L167F 657 ± 3
L167F

**Fig. S6.** The inhibitory activities ($IC_{50}$) of WU-04 (**A-C**) and Ensitrelvir (**D-G**) against 3CLpro were evaluated using a FRET-based assay. The data represent the mean ± SD of three independent measurements. The data of the WT 3CLpro in Figs. S6A and S6D are also used in Figs. 1B and S6E, respectively.

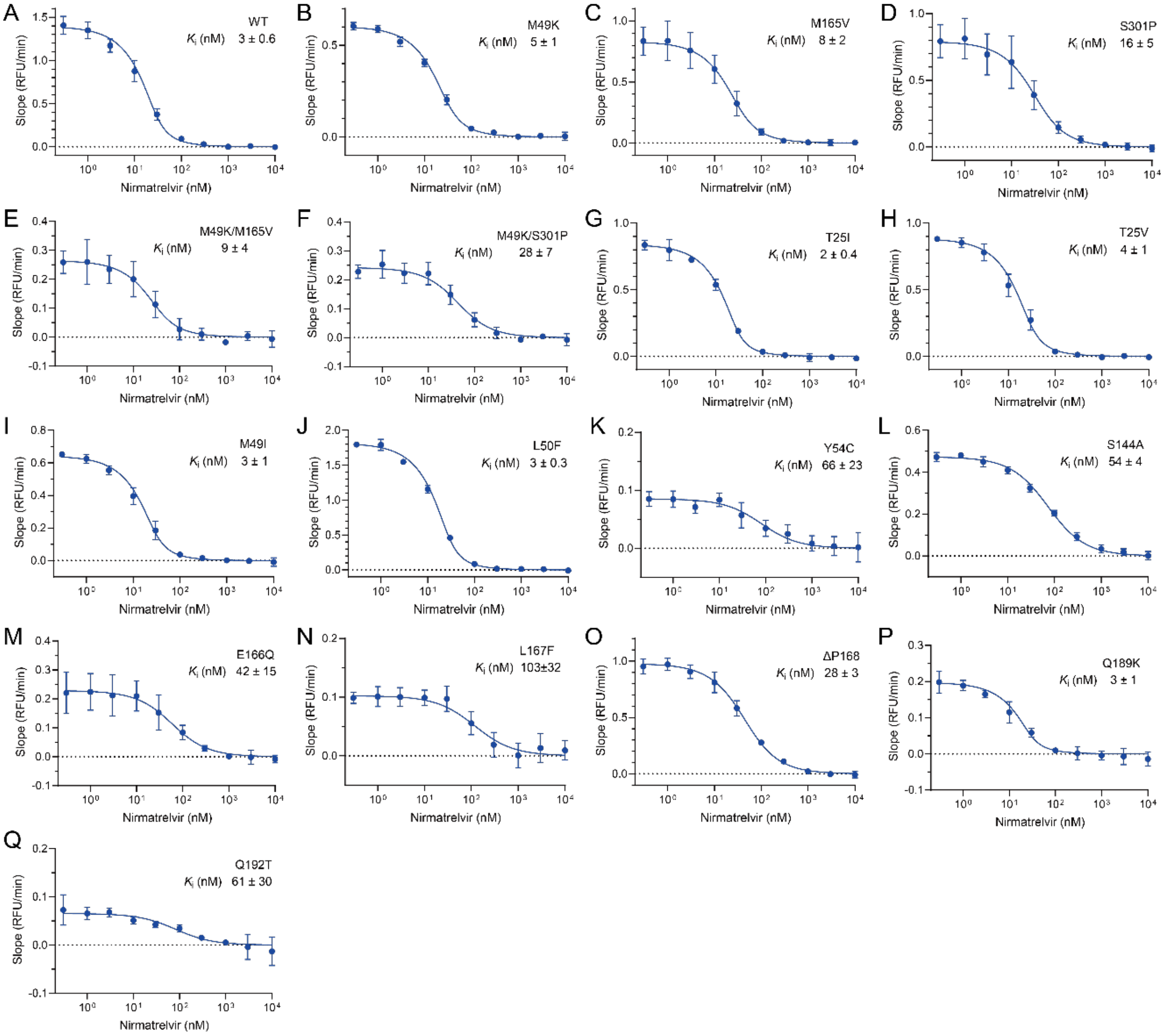


**Fig. S7.** The inhibitory constants ($K_i$) of Nirmatrelvir against the WT 3CLpro (**A**) and the drug-resistant mutants (**B-Q**) were evaluated using a FRET-based assay. The data represent the mean ± SD of three independent measurements.

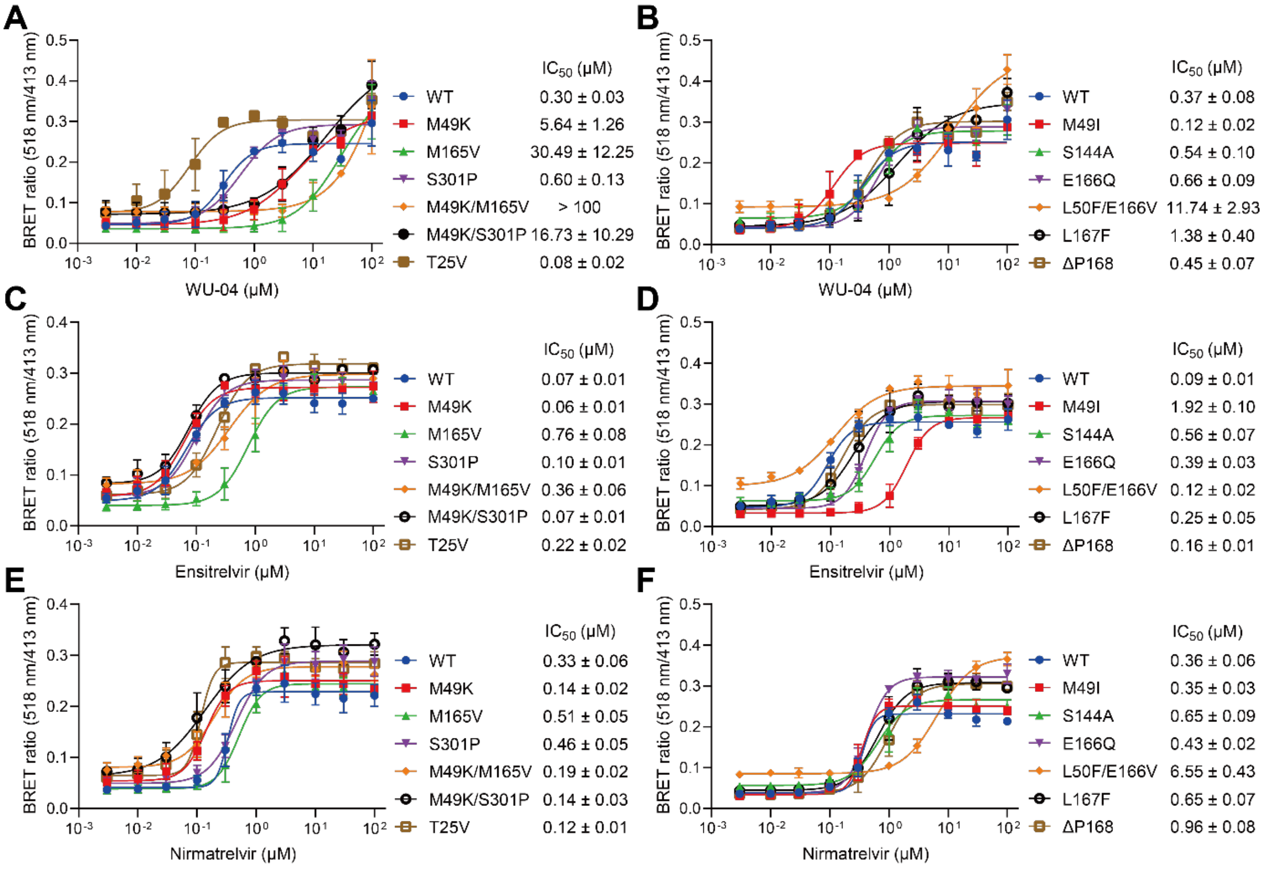


**Fig. S8.** The inhibitory activities of WU-04 (**A**, **B**), ensitrelvir (**C**, **D**) and nirmatrelvir (**E**, **F**) against 3CLpro were evaluated by a BRET-based assay in HEK 293T cells. The data represent the mean ± SD of three independent measurements.

**Table S1.** 3CLpro (NSP5) mutation counts in the GISAID database

| Position | WT (count) | Mutant (count) |
|---|---|---|
| 49 | M (15548971) | K (24); I (2344) |
| 165 | M (15546456) | V (18); Y (4758) |
| 301 | S (15550494) | P (22) |
| 25 | T (15550539) | I (1043); V (1) |
| 50 | L (15546092) | F (5328) |
| 54 | Y (15551450) | C (12) |
| 144 | S (15551564) | A (19) |
| 166 | E (15546519) | Q (4764); V (40) |
| 167 | L (15551296) | F (21) |
| 168 | P (15550992) | deletion (161) |
| 172 | H (15551528) | Y (26) |
| 189 | Q (15550231) | deletion (1192); K (183) |
| 192 | Q (15550004) | deletion (1314); T (221) |

**Table S2.** Data collection and refinement statistics (statistics for the highest-resolution shell are shown in parentheses)

| | *M49K* | *M165V* | *S301P* | *M49K/M165V* | *M49K/S301P* | *M49K/S301P+ WU-04* |
|---|---|---|---|---|---|---|
| *Wavelength* | 1.541 | 1.541 | 1.541 | 1.541 | 1.541 | 1.541 |
| *Resolution range* | 29.66 - 1.5 (1.554 - 1.5) | 28.01 - 2.2 (2.279 - 2.2) | 27.87 - 2.0 (2.072 - 2.0) | 29.63 - 1.5 (1.554 - 1.5) | 29.56 - 2.21 (2.289 - 2.21) | 30 - 1.65 (1.709 - 1.65) |
| *Space group* | I 1 2 1 | I 1 2 1 | I 1 2 1 | I 1 2 1 | P 1 21 1 | I 1 2 1 |
| *Cell dimension a, b, c* (Å) | 51.5257, 81.6172, 90.2145 | 44.6974, 53.6278, 114.183 | 44.6192, 53.4037, 113.384 | 51.582, 80.1127, 90.3169 | 48.8593, 107.074, 54.261 | 51.5706, 81.3544, 89.5668 |
| *Cell dimension α，β，γ (°)* | 90, 96.9412, 90 | 90, 101.126, 90 | 90, 100.529, 90 | 90, 96.8662, 90 | 90, 103.666, 90 | 90, 97.3029, 90 |
| *Asymmetric unit* | 1 | 1 | 1 | 1 | 2 | 1 |
| *Total reflections* | 600407 (37929) | 45420 (4516) | 86547 (8225) | 592321 (36637) | 159440 (16557) | 464381 (28789) |
| *Unique reflections* | 59233 (5920) | 13558 (1331) | 17820 (1750) | 58288 (5781) | 27130 (2692) | 44111 (4387) |
| *Multiplicity* | 10.1 (6.4) | 3.4 (3.4) | 4.9 (4.7) | 10.2 (6.3) | 5.9 (6.2) | 10.5 (6.6) |
| *Completeness (%)* | 99.57 (99.41) | 99.39 (98.95) | 99.65 (99.60) | 99.93 (99.53) | 99.77 (100.00) | 99.98 (100.00) |
| *Mean I/sigma(I)* | 24.16 (1.68) | 10.53 (2.89) | 14.69 (3.85) | 28.44 (1.57) | 16.83 (4.56) | 33.21 (6.49) |
| *Wilson B-factor* | 15.78 | 31.49 | 24.66 | 16.16 | 26.40 | 12.78 |
| *R-merge* | 0.06024 (1.044) | 0.1404 (0.4566) | 0.08929 (0.3017) | 0.04897 (1.12) | 0.1172 (0.3458) | 0.05714 (0.2446) |
| *R-meas* | 0.06296 (1.135) | 0.1661 (0.5359) | 0.1006 (0.3394) | 0.05135 (1.22) | 0.1289 (0.3775) | 0.05971 (0.2658) |
| *R-pim* | 0.0179 (0.436) | 0.08743 (0.2766) | 0.04548 (0.1533) | 0.01508 (0.4758) | 0.05306 (0.1502) | 0.01704 (0.1029) |
| *CC1/2* | 1 (0.869) | 0.968 (0.337) | 0.996 (0.906) | 1 (0.749) | 0.994 (0.914) | 0.999 (0.976) |
| *CC** | 1 (0.964) | 0.992 (0.71) | 0.999 (0.975) | 1 (0.925) | 0.999 (0.977) | 1 (0.994) |
| *Reflections used in refinement* | 59016 (5912) | 13528 (1323) | 17819 (1750) | 58283 (5779) | 27111 (2692) | 44109 (4387) |
| *Reflections used for R-free* | 1993 (199) | 1355 (133) | 1782 (175) | 2000 (198) | 1482 (147) | 1825 (182) |
| *R-work* | 0.1942 (0.2494) | 0.2084 (0.2670) | 0.1792 (0.1995) | 0.1961 (0.2700) | 0.2281 (0.2830) | 0.1626 (0.1607) |
| *R-free* | 0.2053 (0.2942) | 0.2511 (0.3048) | 0.2352 (0.2833) | 0.2084 (0.2703) | 0.2806 (0.3522) | 0.1810 (0.2080) |
| *CC (work)* | 0.963 (0.910) | 0.956 (0.803) | 0.966 (0.922) | 0.964 (0.863) | 0.942 (0.847) | 0.966 (0.950) |
| *CC (free)* | 0.967 (0.815) | 0.944 (0.637) | 0.949 (0.847) | 0.959 (0.854) | 0.890 (0.786) | 0.964 (0.935) |
| *Number of non-hydrogen atoms* | 2548 | 2422 | 2435 | 2529 | 4841 | 2681 |
| *macromolecules* | 2290 | 2367 | 2341 | 2299 | 4740 | 2291 |
| *ligands* | 0 | 0 | 0 | 0 | 0 | 34 |
| *solvent* | 258 | 55 | 94 | 230 | 101 | 356 |
| *Protein residues* | 296 | 306 | 303 | 298 | 612 | 296 |
| *RMS (bonds)* | 0.009 | 0.002 | 0.008 | 0.006 | 0.002 | 0.009 |
| *RMS (angles)* | 1.17 | 0.46 | 0.94 | 0.84 | 0.54 | 1.11 |

| | | | | | | |
|---|---|---|---|---|---|---|
| ***Ramachandran favored (%)*** | 98.63 | 96.71 | 97.67 | 96.94 | 96.88 | 98.63 |
| ***Ramachandran allowed (%)*** | 1.37 | 3.29 | 1.99 | 3.06 | 3.12 | 1.37 |
| ***Ramachandran outliers (%)*** | 0.00 | 0.00 | 0.33 | 0.00 | 0.00 | 0.00 |
| ***Rotamer outliers (%)*** | 0.00 | 0.38 | 0.77 | 0.00 | 0.57 | 0.00 |
| ***Clashscore*** | 1.54 | 1.50 | 2.59 | 1.32 | 3.73 | 1.53 |
| ***Average B-factor*** | 23.95 | 40.10 | 28.87 | 23.27 | 33.23 | 17.79 |
| ***macromolecules*** | 23.11 | 40.19 | 28.81 | 22.48 | 33.34 | 16.35 |
| ***ligands*** | | | | | | 14.19 |
| ***solvent*** | 31.42 | 36.38 | 30.49 | 31.22 | 28.44 | 27.36 |
| ***Number of TLS groups*** | 1 | 1 | 1 | | 1 | 1 |
